# Comprehensive map of ribosomal 2′-O-methylation and C/D box snoRNAs in *Drosophila melanogaster*

**DOI:** 10.1101/2023.05.25.542231

**Authors:** Athena Sklias, Sonia Cruciani, Virginie Marchand, Mariangela Spagnuolo, Guillaume Lavergne, Valérie Bourguignon, René Dreos, Eva Maria Novoa, Yuri Motorin, Jean-Yves Roignant

**Author notes:** Correspondence (J.-Y. Roignant).

## Abstract

During their maturation, ribosomal RNAs (rRNAs) are decorated by hundreds of chemical modifications that participate in proper folding of rRNA secondary structures and therefore in ribosomal function. Along with pseudouridine, methylation of the 2′-hydroxyl ribose moiety (Nm) is the most abundant modification of rRNAs. The majority of Nm modifications in eukaryotes are placed by Fibrillarin, a conserved methyltransferase belonging to a ribonucleoprotein complex guided by C/D box small nucleolar RNAs (C/D box snoRNAs). These modifications impact interactions between rRNAs, tRNAs and mRNAs, and some are known to fine tune translation rates and efficiency. In this study, we built the first comprehensive map of Nm sites in *Drosophila melanogaster* rRNAs using two complementary approaches (RiboMethSeq and Nanopore direct RNA sequencing) and identified their corresponding C/D box snoRNAs by whole-transcriptome sequencing. We *de novo* identified 61 Nm sites, from which 55 are supported by both sequencing methods, we validated the expression of 106 C/D box snoRNAs and we predicted new or alternative rRNA Nm targets for 31 of them. Comparison of methylation level upon different stresses show only slight but specific variations, indicating that this modification is relatively stable in *D. melanogaster*. This study paves the way to investigate the impact of snoRNA-mediated 2′-O-methylation on translation and proteostasis in a whole organism.

## Introduction

In eukaryotic cells, the path leading to a functional ribosome is orchestrated by approximately 80 ribosomal proteins, 4 ribosomal RNAs (rRNAs) and hundreds of non-ribosomal proteins (1). The 28S, 18S and 5.8S rRNAs are transcribed as one single pre-rRNA molecule from a variable number of gene copies that cluster in the nucleoli. Folding and assembly of the mature rRNAs begins as soon as they are processed in a highly timely and complex way. The correct folding of rRNA is a *sine qua non* for the catalytic activity of ribosomes. This process relies on small nucleolar ribonucleoprotein complexes (snoRNPs) that bring distant nucleotides in three-dimensional proximity but also covalently modify rRNAs (2). The most abundant rRNA modifications are 2′-O-methylation (Nm) and pseudouridine (Ψ). The presence of Nm enhances nucleotide stacking and therefore dictates the flexibility of secondary structures of the rRNA, which influences the core function of the ribosome (3). Previous studies reported that both Nm and Ψ modifications concentrate around functionally crucial domains such as the Peptidyl Transferase Centre (PTC), the A- and the P-sites, the Decoding Centre (DC) as well as the inter-subunit bridge (4, 5).

In the case of Nm, the vast majority of site-specific modifications are deposited by a single snoRNP composed of one small nucleolar RNA (snoRNA) of the C/D box family and two copies of four conserved proteins of which Fibrillarin (Fib) carries the catalytic methyltransferase activity (6). Cryo-EM structures have revealed how these proteins form a cavity that bind simultaneously rRNAs and snoRNAs (7). As their name suggests, C/D box snoRNAs are usually short RNAs (<100 nt) that always possess one box C [RUGAUGA] and one box D [CUGA] motif that are separated by one weaker copy of the same motifs, called boxes C’ and D’. The C and D-box sequences, which are located in close proximity to the 5’ and 3’ ends, respectively, interact and form a kink-turn structure that is recognised by the snoRNP complex.

Maturation and assembly of snoRNAs starts as soon as they are spliced out from the introns of other genes where they are usually encoded. According to the strength of the motif, the boxes C’ and D’ located in the middle of snoRNAs will also fold into a kink turn and form a short hairpin. Most importantly, the 8 to 18nt-sequences upstream of the boxes D/D’ can each contain an AntiSense Element (ASE) which is complementary to different rRNA segments. Canonical C/D box snoRNAs follow the D+5 rule, meaning that they guide Nm deposition on the rRNA site that faces the 5th nucleotide upstream of the boxes D/D’ (8). The presence of two ASEs gives the possibility to some C/D box snoRNAs to guide Nm on two distinct targets. In some organisms such as yeast, the same ASE can hybridise to more than one rRNA target, resulting into 42 snoRNAs for a total of 54 rRNA Nm sites (9). On the other hand, in higher eukaryotes such as humans, there are twice more Nm sites in rRNA (n=109), whereas snoRNAs are relatively more numerous than in yeast due to the existence of multiple isoforms (n=130) (10). Despite having identified Nm sites in multiple organisms, the matching between the sites and their corresponding snoRNA has not yet been systematically addressed, and results mainly from predictions.

The alteration of snoRNP members or of single C/D box snoRNAs can lead to a broad spectrum of translational defects. For example, knock-down (KD) of Fib in human cell lines leads to a decrease in rRNA levels and reduced translation efficiency of transcripts that contain IRES-dependent translation initiation sequences (5, 11). Also, mutation of D243 in the catalytic domain of *nop1*, the yeast ortholog of Fib, leads to a significant reduction of ribosome levels and a relative increase in free 40S subunits (4). Knocking out a single C/D box snoRNA is sufficient to shift the codon usage bias of ribosomes (12). Thus, there is increasing evidence that rRNA modifications can fine-tune translation, yet their impact at the physiological level still remains poorly understood.

In *Drosophila melanogaster*, Nm sites have only been predicted based on their complementarity to snoRNA ASEs, which themselves have been partly annotated based on motif predictions from genomic data (13). Various methods have been developed to detect and quantify Nm-modified sites (14, 15). In this study, we use both RiboMethSeq, a nucleotide-resolution sequencing method that employs chemical probing coupled to next generation sequencing, and direct RNA nanopore sequencing (DRS), a long-read sequencing technology that can sequence native RNA molecules, in which modified sites can be identified based on alterations in the current intensity when the RNA molecules are translocated through the nanopores (16). As for C/D box snoRNA quantification, we obtained the full transcriptome of different tissues using TGIRT, a highly processive reverse transcriptase (RT) enzyme that reads through RNA modifications and complex secondary structures. Putting all techniques together, we built a comprehensive map of 61 rRNA Nm sites in *D. melanogaster*, most of which matched with at least one of the 106 expressed C/D box snoRNAs. Our results show that Nm sites on rRNA are relatively stable upon environmental stresses, yet specific changes were observed depending on the type of stress, suggesting specific adaptations. Our work opens the path for investigating the role of individual snoRNAs in multicellular organisms.

## Materials and Methods

### Drosophila stocks and dissection

*D. melanogaster* CantonS reared at 25°C and 65% humidity were used for all *in vivo* assays. Mated females were chosen for all assays as Fibrillarin expression is higher based on modENCODE. Approximately 30 heads were collected for each replicate. For the oxidative stress assay, we extracted RNA from 30 individuals fed for 8 days with 5% Sucrose and 0.1% Nipagin in the control group and additionally 2.5nM Paraquat (Sigma 856177) in the treated group. Flies submitted to nutrient stress were either fed on complete medium (7g of yeast, 10g of sucrose, 2g of agar and 1.5 mL of 10% Nipagin for 100 mL of H2O) or low nutrient medium medium (2.5g of sucrose, 2g of agar and 1.5mL of 10% Nipagin for 100 mL of H2O) for 10 days (17). Flies submitted to heat stress were grown for 8 days at 29°C, 25°C or 18°C on complete medium and in darkness (18).

### Cell culture, RNA interference and transfection

*D. melanogaster* S2R+ cells were grown in Schneider’s medium (Gibco) supplemented with 10% FBS (Sigma) and 1% penicillin–streptomycin (Sigma). For RNA interference (RNAi) experiments, PCR templates for the dsRNA were prepared using T7 megascript Kit (NEB). dsRNA against bacterial β-galactosidase gene (lacZ) was used as a control for all RNA interference (RNAi) experiments (T7-Fib-F: 5’-ACTTCTTACTGCTTGGGCG; T7-Fib-R: 5’-ACCAATGGCGAGAAGATTG; T7-CG8939-F: 5’-GAAAGACGCGCAAGGATAAG; T7-CG8939-R: 5’-TTGTCACGGAAATCATTGGA). S2R+ cells were seeded at the density of 106 cells/ml in serum-free medium and 7.5 μg of dsRNA was added to 106 cells. After 6 h of cell starvation, serum supplemented medium was added to the cells. dsRNA treatment was repeated after 48 and 96 h and cells were collected 24 h after the last treatment. Effectene (Qiagen) was used to transfect vector constructs in all overexpression experiments following the manufacturer’s protocol.

### RNA isolation and RT-PCR

Total RNA from S2R+ cells and Canton-S tissues was isolated with Trizol (Invitrogen) and treated with DNase I treatment (New England Biolabs) for all assays in this study. 250 ng of DNAsed RNA were retro-transcribed using random hexamer primers with the M-MLV-RT enzyme (Promega). cDNA were amplified with the GoTaq qPCR Master Mix (Promega) and the fluorescence was measure on a QuantStudio 6 Flex System (Applied Biosystems). Relative expression was calculated with the ΔΔCt method (qRT-Fib-F: 5’-GCCATTGGTCTCAACGGAG; qRT-Fib-R: 5’-GAGGGAGTGTTCATTGCGC; qRT-CG8939-F: 5’-GCTCAGAGTCATCCGAATCC; qRT-CG8939-R: CGAACCTTCTTGGCATTTGT).

### Detection of 2′-O-methylation by RiboMethSeq (RMS)

RiboMethSeq (RMS) analysis was performed as previously described (19). Briefly, 150 ng of total RNA were subjected to alkaline hydrolysis for 16 min at 96°C followed by ethanol precipitation. The extremities of RNA fragments were end-repaired and converted to libraries using NEBNext□ Small RNA library preparation kit according to the manufacturer’s recommendations. Libraries were quantified and multiplexed and subjected for high-throughput sequencing on a NextSeq2000 instrument with a 50 bp single read mode.

After removal of adapter sequences by trimmomatic v0.39, reads < 40nt were selected and mapped to the *D. melanogaster* rRNA sequences FBtr0346885 (28S), FBtr0346878 (18S) and FBtr0346887 (5.8S) using bowtie2 in End-to-End mode. Reads’ extremities (5’-ends and 3’-ends) were counted and the different RiboMethSeq scores were calculated. A combination of ScoreMean >0.92 and ScoreA >0.5. ScoreC (MethScore) was used for quantification of the methylation level. Conserved Nm sites between species were done based on the alignment of rRNA sequences with Clustal 2.1 (**Supplementary Table 1**) (20).

### Library Preparation for direct RNA Nanopore sequencing (DRS)

Total RNA extracted from untreated and Fib knockdown samples (Fib_KD) in biological duplicates was DNase-treated with Turbo DNase (Thermofisher, AM2238) for 10’ at 37°C. Subsequently, 1000 ng of each sample was polyadenylated with *E. coli* PolyA Polymerase (NEB #M0276L) for 15 min at 37°C. 200 ng of poly(A)-tailed RNA was ligated to pre-annealed custom barcoded adaptors, following previously published protocols (21). Ligated RNA was prepared for direct RNA sequencing using the SQK-RNA002 kit following the ONT Direct RNA Sequencing protocol version DRS_9080_v2_revI_14Aug2019, with minor changes to allow for sample multiplexing. Briefly, for each sample, barcoded oligonucleotides A and B were mixed in annealing buffer (0.01 M Tris-Cl pH 7.5, 0.05M NaCl) to a final concentration of 1.4 µM each in a total volume of 75 µL. The mixture was then incubated at 94°C for 5 minutes and slowly cooled down (-0.1°C/s) to room temperature. Then, 200 ng of total RNA were ligated to the pre-annealed custom RT adaptors (21) using concentrated T4 DNA Ligase (NEB-M0202T). Ligated RNA was reverse transcribed using Maxima H Minus RT (Thermo Scientific, EP0752) at 60°C for 30 min, without the heat inactivation step. The products were purified using 1.8X Agencourt RNAClean XP beads (Fisher Scientific-NC0068576) and washed with 70% freshly prepared ethanol. 50 ng of reverse transcribed RNA from each reaction were pooled together, and the RNA:DNA hybrid was ligated to the RMX adapter. The mix was purified using 1X Agencourt RNAClean XP beads, washing with Wash Buffer (WSB) twice. The sample was then eluted in Elution Buffer (EB) and mixed with RNA Running Buffer (RRB) prior to loading onto a primed R9.4.1 flowcell. The samples were run on MinION sequencing devices.

### Direct RNA sequencing data pre-processing

Raw fast5 files were processed with the Master of Pores pipeline (version 1.5, (22), https://github.com/biocorecrg/master_of_pores). Fast5 files were demultiplexed with DeePlexiCon using default parameters (21) and basecalled with Guppy basecaller v4.0. (https://nanoporetech.com). Reads were mapped to *D. melanogaster* rRNA sequences, obtained from Ensembl: FBtr0346885 (28S), FBtr0346878 (18S), FBtr0346887 (5.8S) and FBtr0086426 (5S). Mapping was performed using minimap2 v2.17 (https://github.com/lh3/minimap2) with “-ax splice -k 14 -uf” options.

### Analysis of RNA modifications in direct RNA sequencing data

Basecalling features (base quality, mismatch frequency, insertion frequency and deletion frequency) of each 5-mer were extracted from 2 replicates of untreated and Fib KD S2R+ reads using *Epinano* (23), version 1.2 (https://github.com/enovoa/EpiNano). Then, each position at the centre of the 5-mer was assigned a score that consisted of the difference between the scaled sum of the aforementioned frequencies in the untreated and in the Fib KD sample, as previously described (24). Then, we calculated the median of scores for each rRNA transcript and each replicate. For *de novo* discovery of Nm sites, nucleotides with a score greater than 3x the median score of all positions in the same transcript in both replicates or 5x the median in one of the replicates were further kept. Given that Nm modifications affects the score over multiple subsequent nucleotides – not just the modified site –, we merged nucleotides passing the previous criterion across 4 consecutive nucleotides into ‘regions’. Regions of ≤6 nts were expanded to 7-mer windows and regions ≥10 nts were split into two 7-mer windows to maximise the occurrence of only 1 Nm site per window. Regions overlapping with the 28S break and 5S rRNA were excluded. For the validation of RMS Nm sites by DRS, we extracted 7-mer windows around each confident RMS site and we kept regions with a score greater than 3x the median score in both replicates or 5x the median in one of the replicates. Integrative Genomics Viewer (IGV) version 2.8.13 was used to visualise mismatches and *Epinano* scores.

### Preparation of TGIRT-Seq libraries

Total RNA from S2R+ cells and Canton-S tissues was isolated with Trizol (Invitrogen) and 3 µg were DNAsed with DNase I treatment (New England Biolabs). RNA was directly ribodepleted with the riboPOOL kit (siTOOLs) that was completed with oligos hybridising to 5S and 2S rRNAs following the manufacturer’s instructions. The quality of RNA and ribodepletion was assessed on an Agilent 2100 Bioanalyzer. The purified ribodepleted RNA samples were fragmented for 3 minutes at 94°C in Magnesium RNA Fragmentation Module (New England Biolabs) and 3’ ends were dephosphorylated with T4 polynucleotide kinase (Lucigen). Between 50 and 75 ng were retrotranscribed in cDNA with 1.5 µL of template-switching TGIRT enzyme. The remaining library was prepared exactly as in Boivin and Deschamps-Francoeur et al (25). After 12-13 cycles of PCR and a 1.4x cleanup with Ampure XP beads (Bekman-Coulter), the profile of TGIRT libraries were evaluated once more by Bioanalyzer (± 250 bp average). Finally, the libraries were sequenced on an Illumina Next-seq 500 platform (2x150) yielding between 9.5 and 19.9 million paired-end reads.

### TGIRT-seq data analysis

Adapters and low quality reads were trimmed from fastq files with cutadapt (version 3.4, (26)). rRNA sequences were filtered out with bowtie2 (-p 20 -L 15 -k 20 --fr --end-to-end) losing up to 4% of reads. The remaining reads were mapped with STAR as described in (27) (version 2.7.8a, (28)). Finally the reads were counted with CoCo (cc -c both -p -s 1), a pipeline that is efficient in attributing reads to genes that are nested in others, such as snoRNAs (29). CPM and TPM were taken from the CoCo output. Analysis of differentially expressed genes was done with DeSeq2 (30) and plots were drawn in R using basic functions and ggplot2 (31) as well as on Powerpoint/Inkscape.

### Screening of canonical C/D box snoRNAs with snoScan

To validate the C/D box features and to predict Nm targets of the 143 snoRNAs (biotype: snoRNA and ncRNA), we used both human yeast and mammalian probabilistic search models of snoScan 1.0 (32). All publications used for mapping snoRNAs in *D. melanogaster* are listed on **Supplementary Table 2**. Briefly, 142 snoRNA fasta files with corrected coordinates were compared against the same rRNA sequences used for the RiboMethSeq. Target predictions of expressed snoRNAs matching experimentally identified Nm sites were considered as highly confident. When the snoRNA was expressed but matched an unknown target or no target at all, it was classified as orphan. The target predictions of snoRNAs for which no expression was detected in the tested tissues were considered as lowly confident.

## Results

### Identification of Nm sites on *Drosophila melanogaster* rRNA

Different rRNA annotations are available in *Drosophila melanogaster* including multiple ones from the Ensembl 103 assembly, one GenBank reference that was used for previous snoRNA predictions, and a PDB reference that was used for 3D representations. To facilitate the navigation between previously predicted and novel identified Nm positions (see below), we aligned annotated rRNA sequences with Sanger-sequenced data from both embryonic cell line S2R+ cells and wildtype Canton-S flies (**Supplementary Table 1** and **Supplementary Figure 1A**). The most similar sequences were FBtr0346885 (28S), FBtr0346878 (18S), FBtr0346887 (5.8S) and FBtr0086426 (5S) from the Ensembl assembly. Consistent with previous observations (33), we confirmed that the 28S rRNA has a ”hidden break” based on the lower read coverage that spans nucleotides 1814 to 1858, but also based on the presence on the electrogram of two RNA peaks just below 2 kb that reflects a mixture of 18S and two similarly sized fragments of 28S (**Supplementary Figure 1B-C**). This break is also present in 5.8S, which is in fact shorter in *D. melanogaster* (123 nts) in comparison to human or yeast (157 and 158 nts, respectively) (34).

To map Nm sites onto the rRNA molecules, we generated RiboMethSeq (RMS) libraries from total RNA extracted from S2R+ cells under control (CTR) and Fibrillarin knockdown (Fib-KD) conditions. It is important to note that only transient and partial KD of Fib can be achieved, since this protein is essential for snoRNP maturation and rRNA processing. The construction of RMS libraries is based on the increased resistance to alkaline pH of the phosphodiester bond between the 2′-O-methylated nucleotide and the one following it. As this method is prone to false positives related to RNA structure, sequence and ligation biases (19), we applied different filters in order to ensure a robust and conservative calling of Nm sites (**Figure 1A**). First, we used a combination of scoreMEAN> 0.92 and scoreA2> 0.5 as in previous studies (35) and identified 183 putative Nm sites by their protection against alkaline cleavage. A MethScore, which is a semi-quantitative parameter that reflects the fraction of sites that are methylated, was then computed for each position. In parallel, we extracted the MethScores from data obtained from *in vitro* transcribed rRNAs (IVTs), which are not methylated and can therefore serve as negative controls. We then filtered candidate Nm sites that had an average MethScore in CTR (n=4) that was higher by 0.05 in comparison to the MethScore observed for IVT rRNAs. This filtering reduced the number of putative Nm sites to 92 (**Figure 1A-B, Supplementary Table 3**). The third filter was based on the significant drop of MethScore that was observed after knocking-down Fib (ΔMethScore ≥ 0.05, p-val <0.05, Wilcoxon test). With this criterion, 55 candidate Nm sites out of 92 previously retained candidates had a significantly lower MethScore in Fib-KD conditions and thus were ranked as highly confident sites (**Figure 1A, 1C** and **Supplementary Table 3**). Interestingly, the differential MethScore was as low as 0.03 for 28S-Am1017 and as high as 0.49 for 18S-Gm475, underlining the very heterogenous response of individual Nm level to the reduction of Fib-KD, probably reflecting differential stability and/or activity of snoRNP complexes. Similar behaviour was also observed for human cell lines upon Fib KD, only a fraction of rRNA sites showed substantial decrease in methylation level (5, 11).

**Figure 1.**
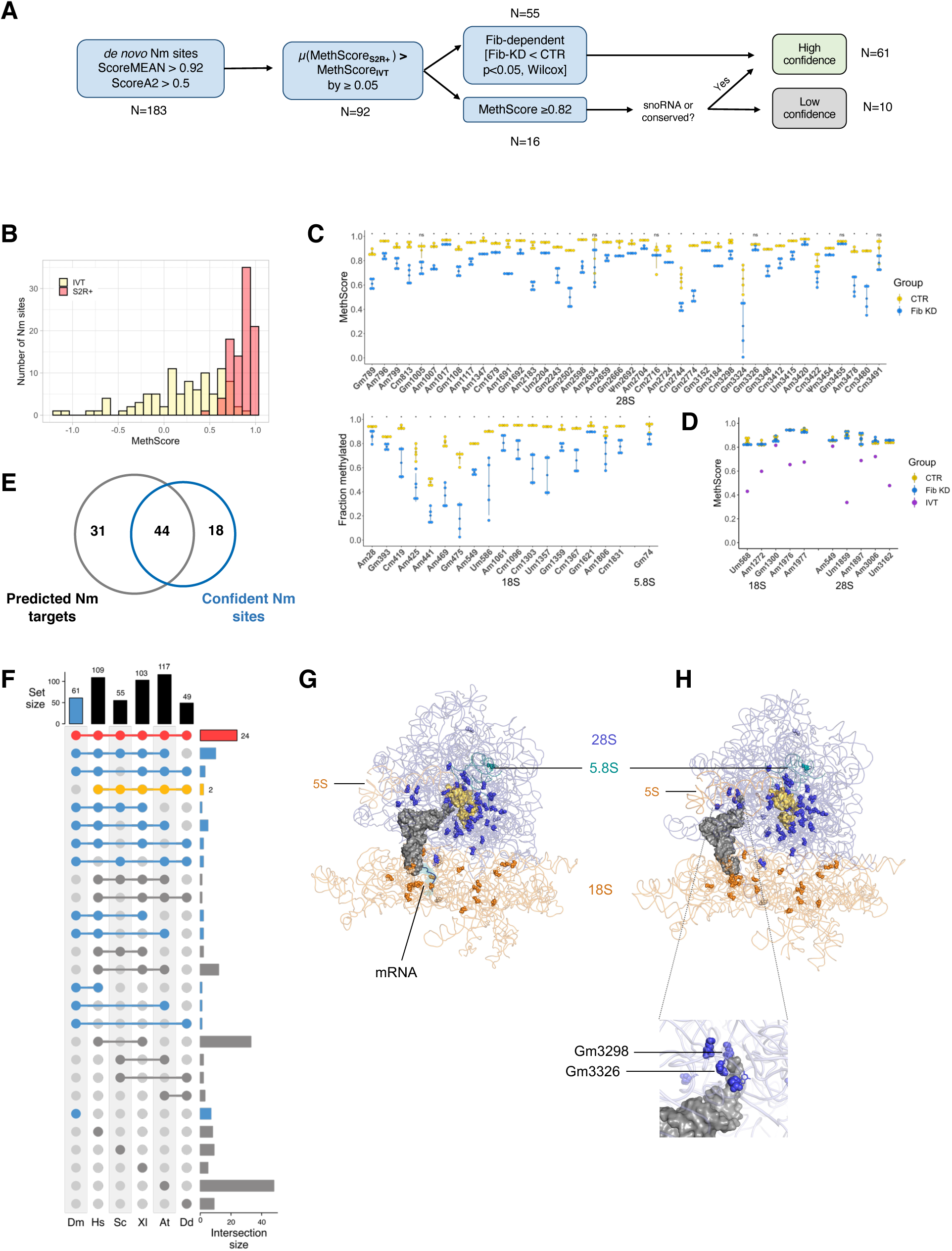
(**A**) Data processing performed for the identification of Nm sites in *D. melanogaster* using RiboMeth-seq. (**B**) Distribution of MethScores for S2R+ cells (red) and of *in vitro* transcribed rRNAs (yellow). (**C**) MethScore for each highly confident Nm site in 28S, 18S and 5.8S rRNAs in control (yellow) and Fibrillarin knockdown (blue) S2R+ cells. Asterisks indicate the significance of loss of methylation (Standard deviation bars, n=4, Wilcoxon test, * p<0.05 and NS p>0.05). (**D**) MethScore for low confidence Nm sites on 18S and 28S rRNAS in control (yellow) or Fibrillarin knockdown (blue) S2R+ cells and in IVT rRNAs (purple). (**E**) Overlap between the 61 confident Nm sites and the Nm candidates predicted from snoRNA complementarity in previous literature (summarised in Supplementary Table 2). (**F**) Upset plot of Nm sites that are conserved across different model organisms (Dm: *D. melanogaster*; Hs: *H.sapiens*; Sc: *S. cerevisiae*; Xl: *X. laevis*; At: *A. thaliana;* Dd: *D. discoideum*). Nm sites found in Dm are highlighted in blue, Nm sites conserved in all species are in red and those that are absent in Dm are highlighted in yellow. (**G, H**) Three-dimensional location of Nm sites in two elongating conformations of ribosomes loaded with a tRNA in the P-site and an mRNA in the decoding centre (testis polysome, 6XU7, reported resolution 4.9 Å) (**G**) and a tRNA in the E-site (embryos ribosome, 4v6w, reported resolution 6 Å) (**H**). The main chain of Nm sites is shown as blue (28S), cyan (5.8S) and orange (18S) spheres and the bases are shown as sticks. The nucleotides facing the Peptidyl Transferase Centre are shown in yellow. Nm sites that are in high proximity with the amino-acid tail of tRNAs are magnified in (**H**).

The remaining population of candidates (37 sites, 92-55=37) did not show Fib-KD dependent reduction in MethScore values (**Figure 1A**). From these, 16 had a MethScore level ≥0.82, a cut-off that was set in a way that the first Nm site above the cut-off was predicted to have a snoRNA match. Since the majority of rRNA Nm methylation events (with the exception of 28S-Gm3455, see below) are snoRNA-dependent in all studied eukaryotic species, the existence of known or predicted C/D-box snoRNA guide complementary to the site can be used as an additional criterium for validation, even in the absence of a clear Fib-dependence. Out of 16 retained sites, 5 had a complementary snoRNA guide and 1 revealed to be highly conserved in other eukaryotic species. These 6 candidates were thus ranked along with the previous high confidence Nm sites (55+5+1 =61). The conserved Nm site with no snoRNA match is 28S-Gm3455 that is predicted to be deposited by CG8939, the orthologue of the stand-alone methyltransferases FTSJ3 in human and Spb1 in yeast (36). Curiously, the knockdown of CG8939 led only to a minor drop of 0.03 at the expected 28S-Gm3455 in comparison to a 0.07 drop at the Nm site located one nucleotide upstream (**Supplementary Figure 2A and 2B**). The ten remaining Nm candidate sites with no associated snoRNA or conservation match were ranked as low confident (**Figure 1D**). For those, the only supporting evidence is high MethScores indicating protection against alkaline cleavage and lower protection observed in IVT. It is not excluded that these could be targets of non-canonical snoRNA guides or yet uncharacterised guide-independent methyltransferases. Alternatively, these low confidence sites could be indicators of other RNA modifications that could contribute to alkaline resistance of RNA. Lastly, biases due to robust secondary structure or in adapter ligation can also create underrepresented fragments in the RMS analysis.

In total, 61 highly confident Nm sites were identified by this successive filtering, with 42 in 28S, 18 in 18S and 1 Nm site in 5.8S, while 5S and 2S rRNAs had no detectable Nm modification (**Table 1**). Only 44 of these highly confident sites overlapped with the 75 Nm sites predicted so far in the literature, essentially based on snoRNA complementarity (**Figure 1E**). We nevertheless checked the MethScore of the remaining 31 predicted sites in the pre-filtered data and none had a value of MethScore above 0.56 or was Fib-dependent. As shown by our expression data (see below), these predicted snoRNAs are not even expressed, suggesting that these Nm sites were inaccurately predicted. Among the 61 highly confident sites, 53 are present in at least one of the five model organisms for which Nm sites have been mapped and 24 are universally conserved in all 5 species (**Figure 1F, Supplementary Table 4**). In contrast, *D. melanogaster* lacks 2 Nm sites that were found to be conserved in all the other studied eukaryotic species.

**Table 1.** : Nm sites in *Drosophila melanogaster* rRNAs identified by RiboMeth-seq

| Target rRNA | Nm site † | Conservation§ | snoRNA (expressed) |
| --- | --- | --- | --- |
| 18S | Am28<sup>N</sup> | DmHYAXDd | FBgn0063388;FBgn0063387;FBgn0063386 |
|  | Gm393<sup>N</sup> | DmHAX | FBgn0065058;FBgn0082931 |
|  | Cm419<sup>N</sup> | DmHYAX | FBgn0086533 |
|  | Am425<sup>N</sup> | DmHYX | FBgn0086056 |
|  | Am441<sup>N</sup> | DmHYAXDd | FBgn0263489 |
|  | Am469<sup>N</sup> | DmHAX | FBgn0086041 |
|  | Gm475<sup>N</sup> | DmDd | FBgn0263851 |
|  | Am549<sup>N</sup> | DmHYAX | FBgn0261973;FBgn0261974 |
|  | Um586<sup>N</sup> | DmHYAXDd | FBgn0060291;FBgn0082926 |
|  | Am1061<sup>N</sup> | DmHYAX | FBgn0082952 |
|  | Cm1096<sup>N</sup> | DmA | FBgn0086055 |
|  | Cm1303<sup>N</sup> | DmHA | FBgn0086061 |
|  | Um1357<sup>N</sup> | DmHYAXDd | FBgn0082949;FBgn0082947;FBgn0082948;FBgn0286758 |
|  | Gm1359<sup>N</sup> | DmHYAXDd | FBgn0086053;FBgn0086054;FBgn0286758 |
|  | Cm1367<sup>N</sup> | Dm | FBgn0082951 |
|  | Gm1621<sup>N</sup> | DmHYAXDd | FBgn0063389 |
|  | Am1806<sup>N</sup> | DmHX | FBgn0086074 |
|  | Cm1831<sup>N</sup> | DmHYAXDd | FBgn0065053 |
| 28S | Gm789<sup>N</sup> | DmH | FBgn0020518 |
|  | Am796<sup>N</sup> | DmHXDd | FBgn0086067 |
|  | Am799<sup>N</sup> | DmHYAXDd | FBgn0086078;FBgn0086079;FBgn0086067 |
|  | Cm813<sup>N</sup> | DmHYAX | FBgn0086062;FBgn0086063 |
|  | Gm1005<sup>N</sup> | DmHYAX | FBgn0063392;FBgn0065047;FBgn0065048;FBgn0086042 |
|  | Am1007<sup>N</sup> | DmHYAX | FBgn0082943;FBgn0082942 |
|  | Am1017<sup>N</sup> | DmHYAX | FBgn0082941 |
|  | Gm1108<sup>N</sup> | DmHYAX | FBgn0082935;FBgn0082934;FBgn0082933;FBgn0082932 |
|  | Am1117<sup>N</sup> | Dm | FBgn0082935;FBgn0082934;FBgn0082933;FBgn0082932 |
|  | Am1347<sup>N</sup> | DmHYAXDd | FBgn0015543 |
|  | Cm1679<sup>N</sup> | DmHYAXDd | FBgn0086068 |
|  | Am1691<sup>N</sup> | DmHYAXDd | FBgn0063377;FBgn0063376;FBgn0086068 |
|  | Gm1692<sup>N</sup> | DmHYAXDd | FBgn0063377;FBgn0063376 |
|  | Am2183<sup>N</sup> | DmHAX | FBgn0086039 |
|  | Um2204<sup>N</sup> | DmHYAXDd | FBgn0260002;FBgn0086051 |
|  | Gm2243<sup>N</sup> | DmHX | FBgn0086028 |
|  | Gm2502<sup>N</sup> | DmHA | snoRNA not found |
|  | Am2598<sup>N</sup> | DmHYAX | FBgn0065071 |
|  | Am2634 | DmHYAXDd | FBgn0065066 |
|  | Am2659<sup>N</sup> | DmHYAXDd | FBgn0086023;FBgn0086024;FBgn0086025 |
|  | Gm2666<sup>N</sup> | DmHYAXDd | FBgn0025881 |
|  | Ψm2692<sup>N</sup> | DmHXDd | FBgn0086603 |
|  | Am2704<sup>N</sup> | DmHAXDd | FBgn0082944;FBgn0082945;FBgn0082946 |
|  | Cm2716 | DmHYAXDd | FBgn0082938;FBgn0082937;FBgn0082936 |
|  | Am2724<sup>N</sup> | Dm | FBgn0082938;FBgn0082937;FBgn0082936 |
|  | Cm2744<sup>N</sup> | DmHAX | FBgn0065073;FBgn0263467 |
|  | Gm2774<sup>N</sup> | DmHYAXDd | FBgn0082931;FBgn0082930;FBgn0082929 |
|  | Gm3152<sup>N</sup> | DmHYAXDd | FBgn0063381;FBgn0063380;FBgn0063379;FBgn0063378 |
|  | Gm3184 | DmHAXDd | FBgn0086064;FBgn0086065 |
|  | Cm3298<sup>N</sup> | Dm | FBgn0082940;FBgn0082939 |
|  | Gm3324<sup>N</sup> | DmYADd | FBgn0086075 |
|  | Gm3326<sup>N</sup> | DmHYAX | FBgn0082928;FBgn0082927;FBgn0086075 |
|  | Gm3348<sup>N</sup> | DmHYAXDd | FBgn0063375;FBgn0063374;FBgn0063373 |
|  | Cm3412<sup>N</sup> | DmHAXDd | FBgn0060292;FBgn0060291;FBgn0065070 |
|  | Um3415<sup>N</sup> | Dm | FBgn0060292;FBgn0060291;FBgn0086048;FBgn0086049 |
|  | Am3420<sup>N</sup> | Dm | FBgn0086057 |
|  | Cm3422<sup>N</sup> | Dm | FBgn0086069 |
|  | Ψm3454 | DmHYAXDd | snoRNA not found (CG8939-dependent) |
|  | Gm3455 | DmHYAXDd | snoRNA not found (SPB1 in yeast) |
|  | Am3478<sup>N</sup> | DmHYAXDd | FBgn0063385;FBgn0063384;FBgn0063383;FBgn0063382 |
|  | Cm3480<sup>N</sup> | DmYADd | FBgn0063385;FBgn0063384;FBgn0063383;FBgn0063382 |
|  | Cm3491<sup>N</sup> | DmHYAXDd | FBgn0083988;FBgn0083989;FBgn0086022 |
| 5.8S | Gm74 | DmHAX | FBgn0086066 |

† : Ensembl 104 28S/FBtr0346885; 18S/FBtr0346878; 5.8S FBtr0346887
§ : Dm: *D. melanogaster*; Hs: *H. sapiens*; Sc: *S. cerevisiae*; Xl: *X. laevis*; At: *A. thaliana*; Dd: *D. discoideum*<sup>N</sup> : Nm site validated with Nanopore

Consistent with previous work, we found that most Nm sites in rRNA were highly methylated (54/61) (**Figure 1C**, **Table 1**). The 8 Nm sites that were partially methylated (MethScore <0.82) were still Fib-dependent (**Figure 1C**). Intriguingly, five of them were located back-to-back from position 425 to 549 in the 18S rRNA. Cryo-EM structures revealed that these sites are less protected by ribosomal proteins (37, 38). In line with observations for other organisms, Nm sites on the large ribosomal subunit (LSU) are mainly clustered in the immediate surrounding of the PTC and the peptide exit tunnel (PET) (**Figure 1G-H** and **Supplementary Figure 3A** for 2D-projection). In addition, about 1/3 of 18S Nm modifications are gathered around the decoding centre (DC) while the other 2/3 are distributed across the 18S rRNA (**Supplementary Figure 3B** for 2D-projection). As for the only Nm site of 5.8S rRNA, it is found on the wall of the PET. The existence of cryo-EM snapshots containing a tRNA in the E site (38) (**Figure 1H**) revealed a so far undescribed cluster of 4 Nm sites on the 28S rRNA that are in proximity to the amino acid attachment site of tRNAs – two of them, Gm3326 and Cm3298 being as close as 3.5 Å to the closest tRNA atom. The examination of the Nm sites in the context of the secondary structure showed that they are primarily located at the base of helices and in small bulges or loops and are systematically absent from expansion segments, which are usually species-specific (**Supplementary Figure 3A and 3B**).

### Orthogonal validation of Nm sites using nanopore sequencing

Since the presence of other modifications (e.g., pseudouridine) as well as the secondary structure of rRNAs can occult the detection of Nm sites and/or lead to false positives candidates in RMS analysis (19) we sought to use an alternative approach to identify and validate RMS-identified Nm sites. Specifically, we employed direct RNA nanopore sequencing (DRS), which has been shown to detect Nm sites on the basis of an increased base-calling error in the 5-mer window containing each Nm-modified site (24).

Briefly, two independent replicates of Fib-KD and two CTR samples that were used for RMS were also sequenced using DRS. The *de novo* Nm discovery approach (described in the Methods section) identified 53 regions (7-mers) as significantly altered when comparing Fib-KD and CTR samples. Of these regions, 40 overlapped with a total of 42 Fib-dependent and 3 non-dependent Nm sites identified using RMS (**Figure 2A**, **Table 1**). Nevertheless, some of them were part of a region overlapping with two RMS sites, suggesting that the signal from a single Nm site can mislead the detection of a neighbouring Nm site, due to the spreading of the Nm signal across several nucleotides (24). On a similar note, the longer the ‘regions’ were before re-sizing, the more likely it was that they overlapped with two consecutive Nm sites (**Supplementary Figure 4A**). For example, our *de novo* Nm detection pipeline could not identify that a given region contained 2 modified sites in the following cases: 28S-Am3478 and Cm 3480, or Gm3324 and Gm3326, which are in short distance from one another (**Figure 2B**).

**Figure 2.**
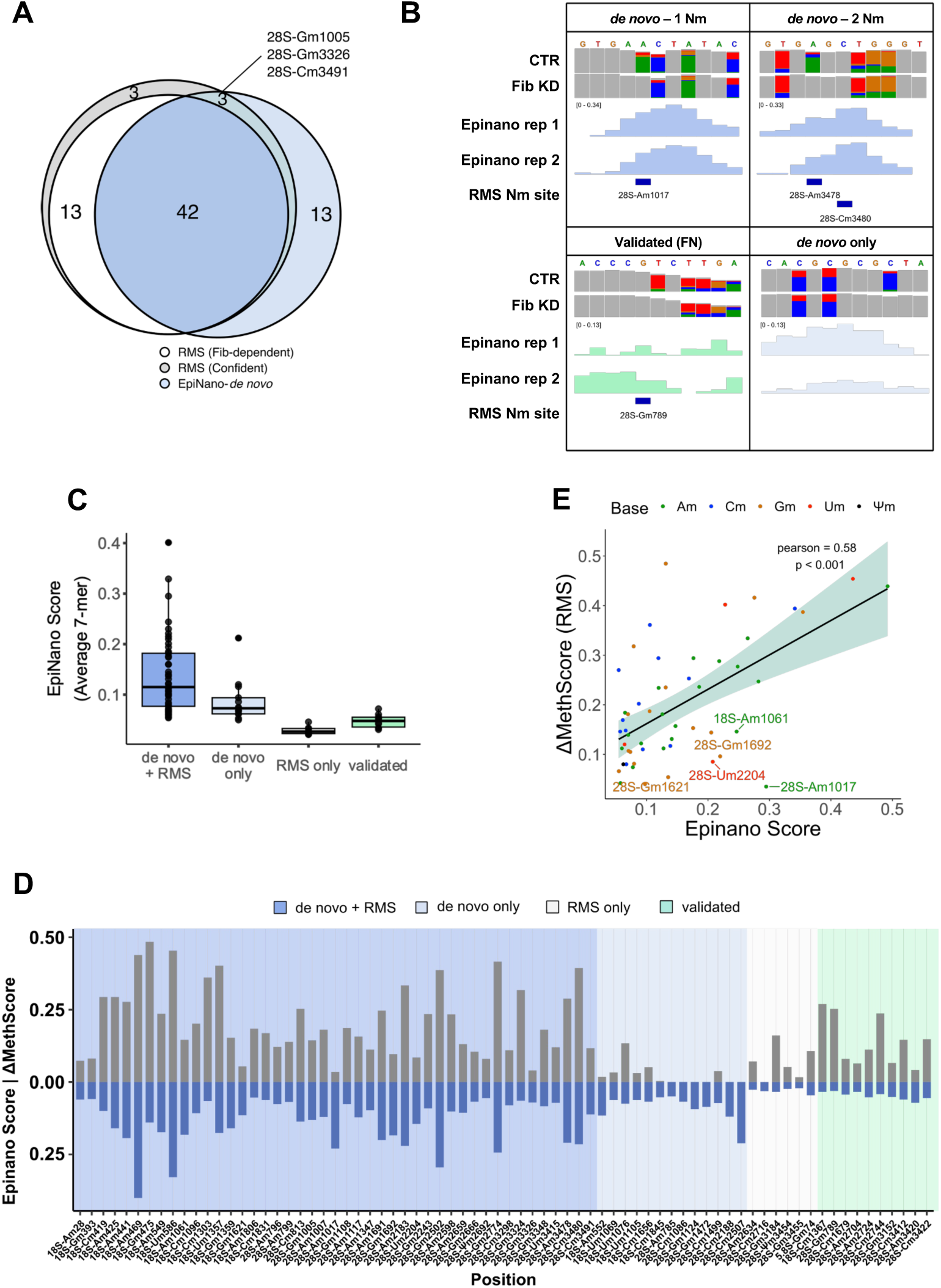
(**A**) High confidence RMS Nm sites (grey), RMS Fibrillarin-dependent Nm sites (white, ΔmethScore ≥ 0.05) that overlap *de novo* nanopore sites (blue). (**B**) IGV tracks of representative examples of *de novo* and validation analysis by DRS. Positions with base-calling allele error frequencies greater than 20% are coloured; grey represents match to reference. (**C**) Distribution of *Epinano* score of 7-mer regions based on whether they were discovered *de novo* or validated and whether they overlapped with RMS Nm sites. (**D**) Differential methylation between Fib KD and CTR S2R+ cells obtained with RiboMethSeq (ΔMethScore, n=4, blue) and with Nanopore (*Epinano* Score, n=2, turquoise) for high confidence Nm sites (n=61). (**E**) Pearson correlation of ΔMethScore and *Epinano* Score for overlapping Nm sites. Nm sites are coloured by base and those with significantly higher *Epinano* Score are labelled with their coordinates.

The 13 regions that were discovered by the *de novo* DRS but not the RMS analysis were in average shorter and had lower *Epinano* score in comparison to overlapping regions (**Figure 2A-C**) (23). We therefore used different approaches to evaluate their validity. Five sites were eliminated as they were also picked in two independent negative controls (CG8939 and CG11447 KDs, data not shown). Three other regions were also discarded as they were generated artificially by splitting one large region that overlapped a single RMS site. In total, only 5 regions remained after this additional filtering, without an obvious explanation, as none of them overlapped with Nm sites found in other organisms (**Supplementary Figure 4B**) and no matching snoRNA could be predicted. They could result from the alteration of the current provoked by other factors such as 2D rRNA structure that can be indirectly affected by distal Nm sites (although the rRNA molecules were linearized during the library preparation to minimize the effect of RNA structure in the nanopore signal). Further work will be necessary to determine whether these regions do indeed encompass real Nm sites.

Finally, we examined the capacity of DRS to validate known RMS sites. For this, 7-mer windows around each confident RMS site were extracted and kept when the *Epinano* score was 3-fold greater than the background (see *Methods*). This way, 10 Nm sites could be validated along with the 45 *de novo* ones, corresponding to a total of 55 Fib-dependent RMS sites (95%) (**Figure 2C** and **D**, **Table 1**). We noted that the sites captured in the validation, but not in the *de novo* approach, were often missed because of the score being altered in shifted windows, or because the score was extremely low in only one of the two replicates, but not the other one as it was the case for 28S-Gm789 (**Figure 2B** and **Supplementary Table 5**). Finally, as expected, the 3 Nm sites that were Fib-independent in the RMS dataset and missed by the *de novo* approach were also missed by the validation approach (**Figure 2A**). Overall, RMS ΔMethScores and *Epinano* scores correlated significantly (Pearson r=0.58, p<0.001), although the amplitude of RMS ΔMethScores was larger than that of *Epinano* scores (**Figure 2D and 2E**). We further noticed that nanopore was more sensitive to differential methylation on Am sites in comparison to RMS, which had no visible bias towards any base (**Figure 2E, Supplementary Figure 4C and 4D**). In conclusion, while 77% of RMS-identified Nm sites were *de novo* discovered with nanopore sequencing, 95% of Fib-dependent RMS-identified sites could be validated with nanopore, indicating that the nanopore approach is a *bona fide* validation tool of Nm modifications on rRNA.

### Identification of C/D box snoRNAs and prediction of Nm targets

A range of expression and genomic data from previous work has led to the identification of 143 transcripts annotated as C/D box snoRNAs in *D. melanogaster* (**Supplementary Table 2** and **Supplementary Figure 5**). To validate snoRNA expression in a comprehensive manner, we sequenced the whole transcriptome of S2R+ cells using the TGIRT-seq approach – a rRNA-depleted library preparation that covers all RNA species (**Supplementary Figure 6A**) (25, 27). Canonical snoRNAs have a characteristic “square” coverage as end-to-end reads span the snoRNA gene and are blunted sharply at the transcript extremities (**Supplementary Figure 6B**). We noticed a discrepancy between coverage and coordinates reaching up to 63 nucleotides (**Supplementary Figure 6B**). Therefore, we redefined new genomic coordinates for the C/D box snoRNAs based on the read coverage (**Table 2**).

**Table 2.**
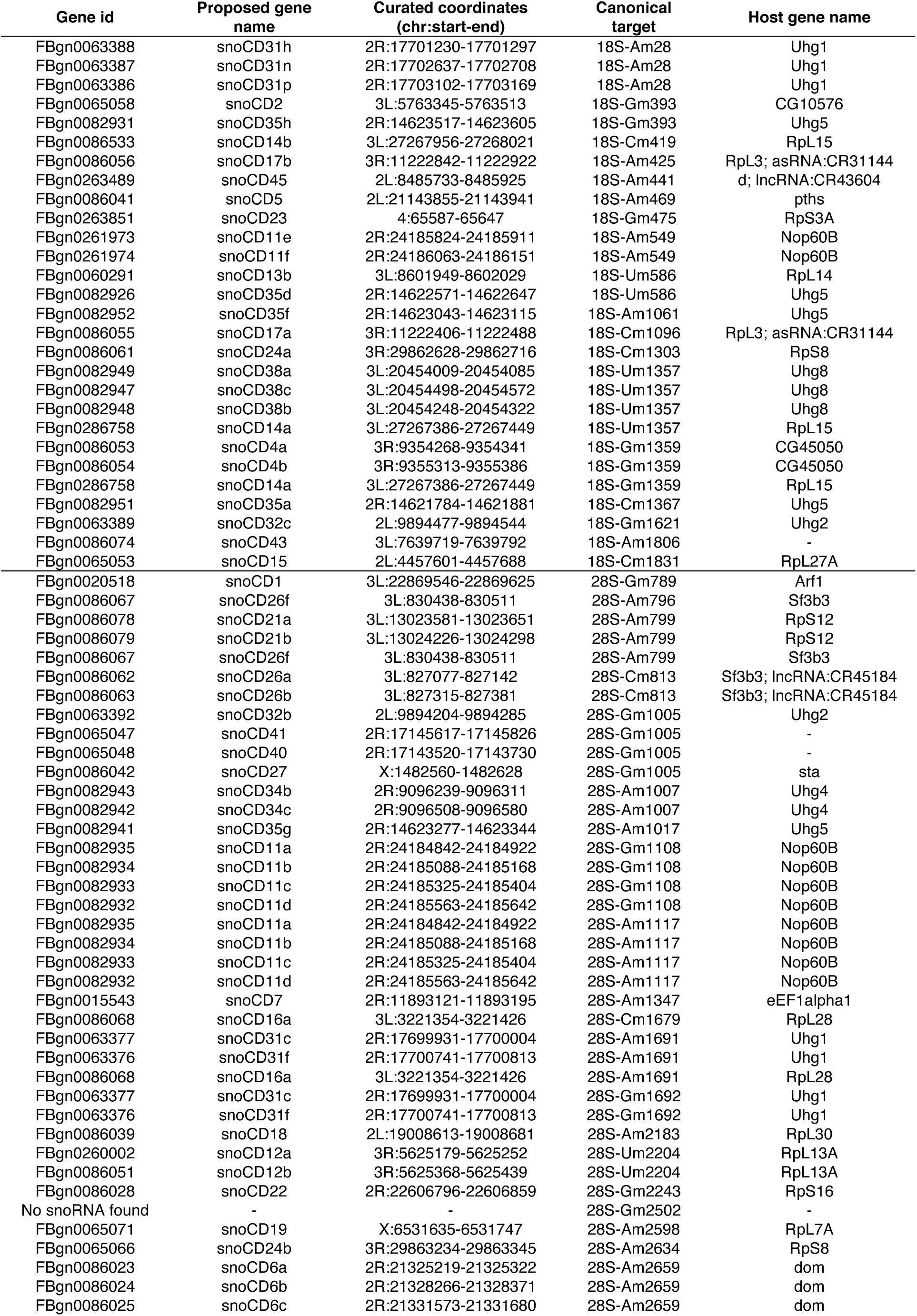
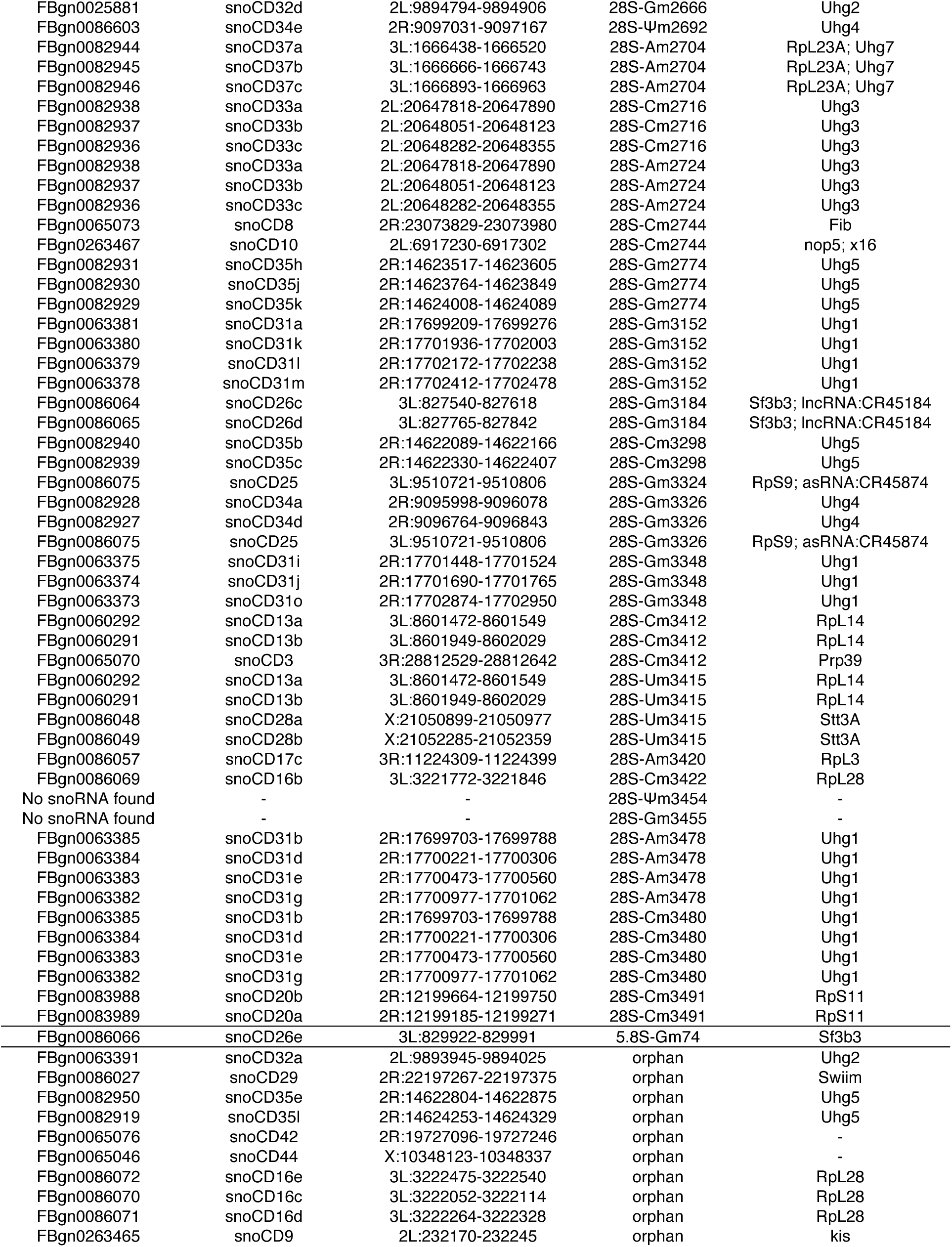
Expressed C/D box snoRNAs and their predicted Nm sites

In total, we found 105 out of the previously annotated 143 snoRNAs expressed in S2R+ cells (**Figure 3A, first and middle panel**) and we identified one novel snoRNA candidate (see below). Interestingly, about two-thirds of the non-expressed snoRNAs were predicted from genomic sequences without any expression evidence in literature nor in any stage or tissue in modENCODE datasets, suggesting that they might have been wrongly annotated (**Figure 3A, Supplementary Figure 6C, Supplementary Table 6**). To search for possible additional expressed snoRNA we repeated the TGIRT-seq experiments with samples derived from other *D. melanogaster* tissues, including heads and ovaries (data not shown). While we found variation in snoRNA expression among different tissues, we did not detect any additional C/D box snoRNA beyond the 106 already detected in S2R+ cells. This indicates that this set of snoRNAs is the most confident one while the predicted snoRNAs that are not detected are either non-existing or expressed in a very localized manner.

**Figure 3.**
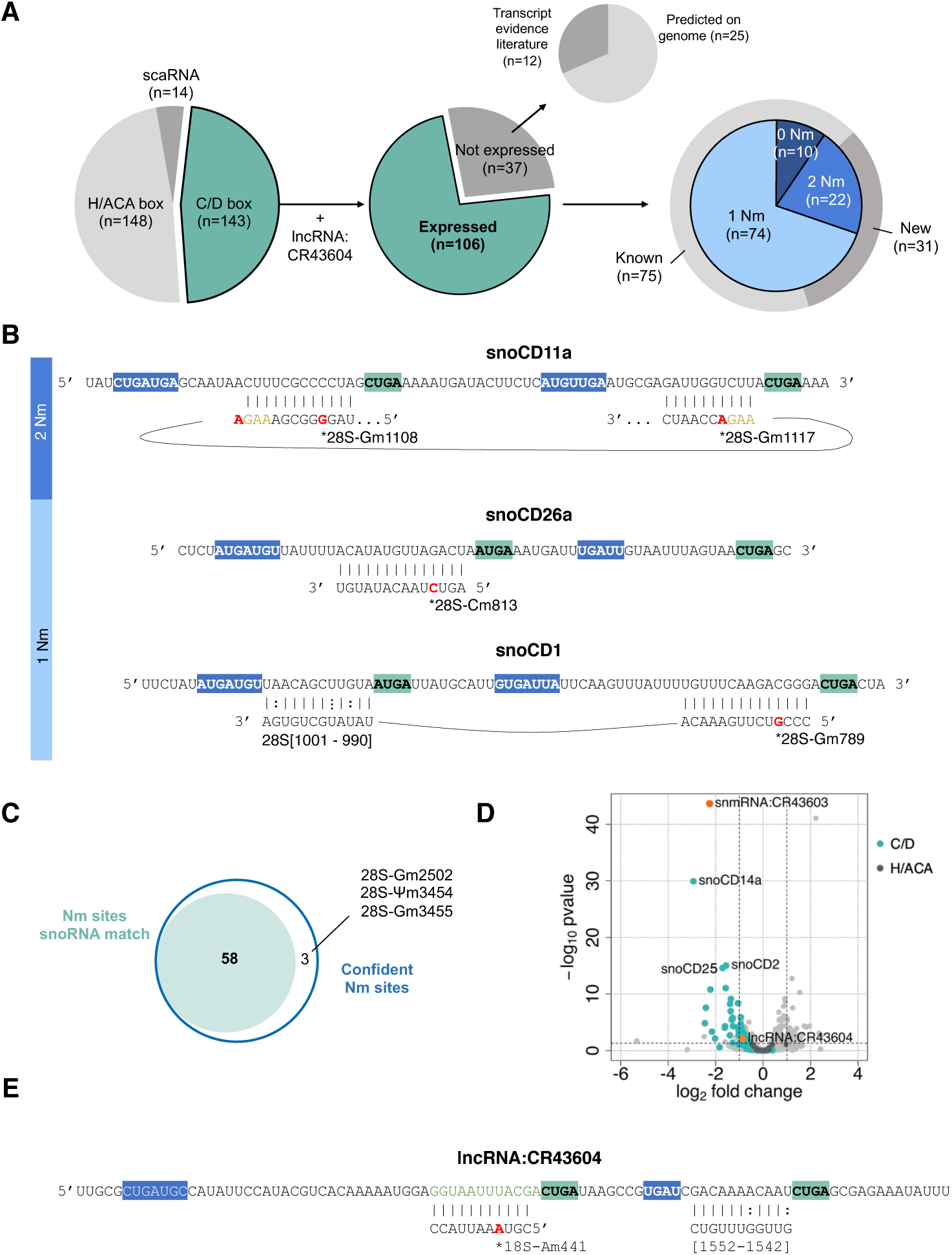
(**A**) Distribution of transcripts labelled with ”biotype:snoRNA” in the annotation BDGP 6.32 (left pie chart). Number of C/D box snoRNAs that are expressed in S2R+ cells (middle pie chart). The upper pie chart indicates whether the non-expressed snoRNAs have previously been annotated based on expression evidence or based on predictions from genomic sequences. The right pie chart enumerates in the first layer the snoRNAs matching a 1, 2 or no Nm sites (light blue to dark blue) and in the second layer are enumerated the C/D box with a previously known or a new rRNA target prediction. (B) Examples of C/D box snoRNAs with 2 or 1 ASE and their predicted interactions with rRNAs. C/C’ boxes are highlighted in blue and D/D’ boxes are highlighted in turquoise. (C) High confident Nm sites with a snoRNA match. (D) Volcano plot of whole-transcriptome changes following Fib KD. C/D box and H/ACA snoRNAs are highlighted in turquoise and dark grey respectively. The horizontal and vertical dashed lines illustrate respectively the FDR =0.05 and |logFC|>1. Two Fib-dependent lncRNAs are highlighted in orange (E) Predicted snoRNA features of lncRNA:CR43604 and interaction with 18S-Am441.

The next step was to match each snoRNA to a target nucleotide in rRNAs. For this purpose various prediction tools have been developed over the years such as snoScan (32). Briefly, we favoured the prediction with the higher score (detailed pipeline in **Supplementary Figure 5** and **Supplementary Methods**). When multiple rRNA positions were predicted for a given snoRNA, the secondary prediction was also kept if it matched a consecutive confident Nm site. As a result, we *de novo* predicted that most of the expressed C/D box snoRNAs (96/106) can target at least one confident Nm site. More specifically, 74 matched to only one confident Nm site, 10 remained unmatched (“orphan”) while 22 snoRNAs were predicted to target at least 2 Nm sites on rRNAs (**Figure 3A, right panel**). An example is shown in **Figure 3B** where consecutive Nm sites can be targeted by the same snoRNA (**Figure 3B**, **Table 2**). Although it is unknown whether these are deposited in parallel or sequentially, we could speculate that the deposition of 28S-Gm1108, Am1117 and also perhaps Am1017 “seals” the conformation of helixes 33 and 35a. Yet, the majority of snoRNAs correspond to a single rRNA Nm site e.g. snoCD26a, although it is not rare that a second ASE matches to rRNA sequences without leading to the deposition of Nm (**Figure 3B**). As already demonstrated in other species, snoRNA:rRNA interaction contributes to the correct folding of rRNAs during ribosome biogenesis. In sum, the combination of the RMS and of TGIRT-seq data led to the prediction of 31 new snoRNA:rRNA matches (**Figure 3A, right panel, Supplementary Table 6**) as well as the attribution of at least one snoRNA to 58 out of the 61 Nm sites (**Figure 3C**).

In line with previous work, we found that the depletion of Fib led to a significant downregulation of 40 C/D box snoRNAs (FDR<0.05) and to a visible downregulation of most of the remaining expressed C/D box snoRNA transcripts (**Figure 3D, Supplementary Figure 6D**). In contrast, H/ACA snoRNAs that are often co-expressed in neighbouring introns, remained unaffected and are globally more highly expressed than C/D box snoRNAs (**Figure 3D, Supplementary Figure 6D**). Interestingly, two uncharacterised ncRNAs, snmRNA:CR43603 and lncRNA:CR43604, were also downregulated. By screening for snoRNA features in these downregulated genes, we found that the lncRNA CR43604 is predicted to target 18S-Am441, a Nm site that is conserved and partially methylated in all other model organisms, for which no canonical snoRNA was predicted so far (included in **Figure 3E**). This lncRNA is covered by two distinct blocks of reads. The C/D boxes and the ASE are identified in the second block.

### Novel nomenclature for C/D box snoRNAs

As it stands, the existing nomenclature for about two-thirds of C/D box snoRNAs in *D. melanogaster* contains the information of the predicted rRNA target (e.g. snoRNA:Me28S-G3113a). We found that this annotation could not only be misleading but it was also created using obsolete rRNA references. First our results demonstrated that several predicted targets were actually false while others were missing. Second, we found that a snoRNA can guide Nm at different rRNA sites, which is not reflected in the currently existing snoRNA annotation. Third there is growing evidence that snoRNAs can target other RNA species in addition to rRNAs, and therefore the current annotation is clearly too conservative (39). Inspired from the human nomenclature (SNORDxx) and following the guidelines of the HUG Gene Nomenclature Committee, we propose a new nomenclature that would not mention the nucleotide target, as it is often incorrect or non-exhaustive (40). In addition, we did not wish to pick the same numbering for snoRNAs that are conserved in other species as there is no strict 1-to-1 match for any species. Therefore, we propose to use the acronym “snoCD” followed by a number that is the same in the case of snoRNAs that are encoded in the same host gene but followed by a letter indicating the order of appearance on the coding strand (**Table 2**). Numbering was done arbitrarily following the alphabetical order of host gene names in the Ensembl assembly 104. Importantly, if new C/D box snoRNAs are discovered in the future, the nomenclature could be expanded without altering the present annotations.

As an example, transcript FBtr0113532 that was previously called snoRNA:Me28S-G1083b, can guide Fib on both 28S-G1108 and A1117 as per the validated 28S rRNA reference. It is renamed as snoCD11b, where “11” signifies the alphabetical ranking of the host transcript Nop60B and “b” signifies the snoRNA is second in terms of genomic coordinates. Finally, snoRNAs predicted, but not expressed in our samples, or only predicted from genomic sequences were not renamed.

### rRNA Nm levels are relatively stable during stress conditions

RNA modifications can be dynamically regulated upon different stress and developmental conditions. Although several reports have shown that rRNA Nm modifications are variable in different cell types and developmental stages, no study has quantified Nm levels in response to long environmental stresses (10, 41, 42). Having built a list of confident Nm sites we therefore sought to explore whether this variability can be observed in *D. melanogaster* subjected to different stresses. We selected environmental stresses that affect ribosome composition or abundance and we first challenged female flies to paraquat, a well-known herbicide inducing oxidative stress in cells (43). Flies were fed on medium containing paraquat up to 8 days, to allow enough time for turnover of existing ribosomes (BioNumbers 108025, 108023, 110053) (44). Total RNA from heads was isolated and submitted to RMS. We first noticed that all Nm sites identified in S2R+ cells were found in wildtype Canton-S flies. Overall, we found very little variation on Nm levels in paraquat treated flies versus control conditions (**Figure 4A**) as almost no Nm site varied more than 0.02 in MethScore value. Only Ψm3454 in 28S rRNA showed a mild reduction of 0.05 in MethScore. Thus, we concluded from this experiment that oxidative stress conditions have negligible impact on the establishment and maintenance of Nm on rRNA in *D. melanogaster*.

**Figure 4.**
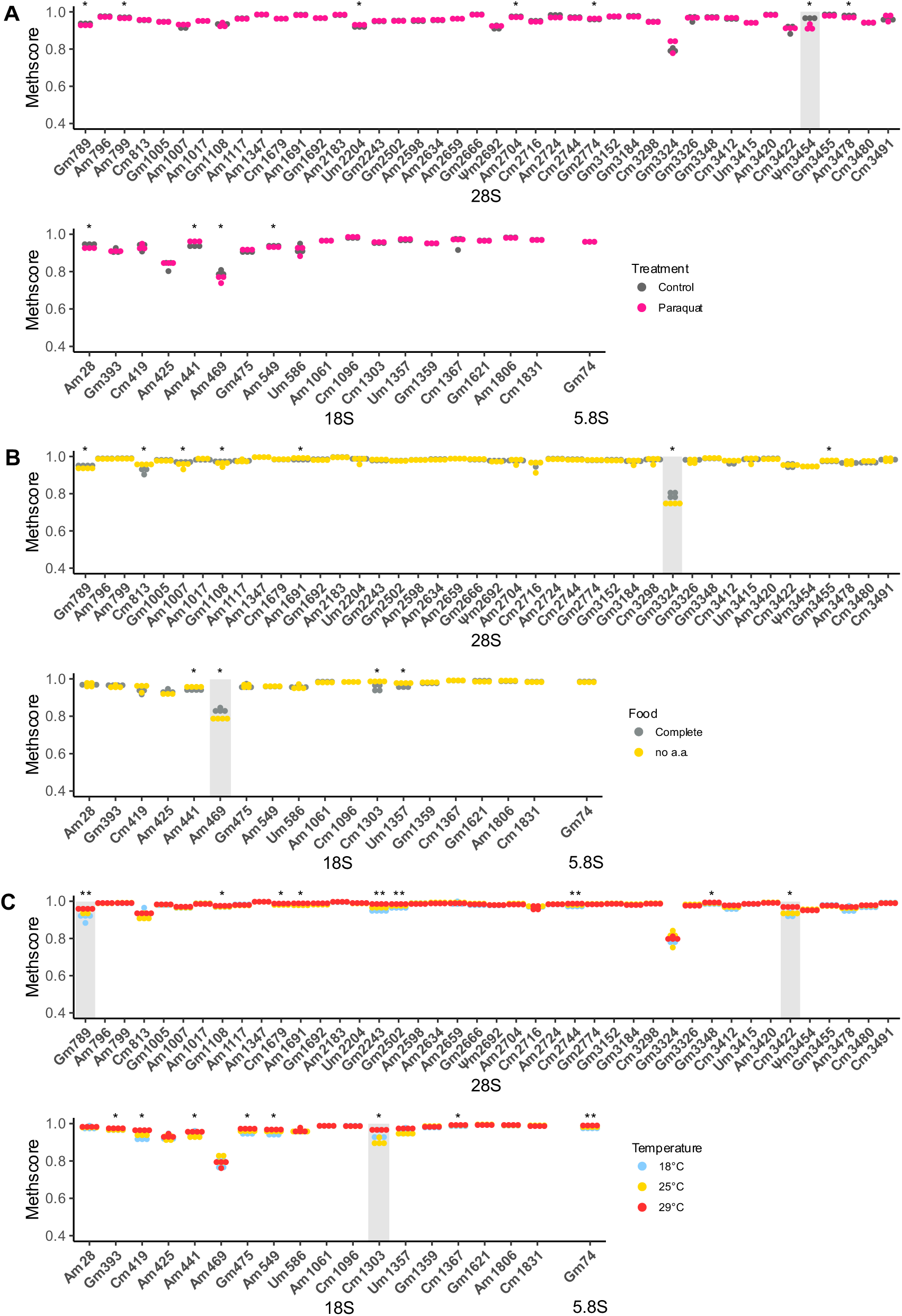
Nm levels in rRNAs from heads of female Canton-S individuals that were exposed (**A**) to paraquat over 8 days (n=3); (**B**) that were fed with complete food medium or amino-acid-depleted and low sugar diet over 10 days (n=4) and (**C**) that were exposed to 18°C, 25°C or 29°C over 8 days (n=4). Asterisks indicate the significance of methylation changes for each Nm site (Kruskal Wallis test, ** p<0.01, * p<0.05 and NS p>0.05). Nm sites with an average ΔMethscore ≥0.05 are highlighted in grey.

We next wondered whether this absence of variability also holds true during other stress conditions. We fed adult females on low nutrient food (no amino acid and 0.25X sugar) for 10 days and monitored the Nm levels on total RNA isolated from heads (17). Again, Nm levels were very stable, despite numerous sites showing a significant difference in nutrient-deprived flies in comparison to controls (**Figure 4B**). Nevertheless, two sites 28S-Gm3324 and 18S-Am469 were both hypomethylated by 0.05 in the nutrient-deprived group. Lastly, we challenged the flies to different temperatures and repeated the RMS assay. This time, 18 sites showed differential Nm methylation (**Figure 4C**). While this number was higher compared to the other stresses, the extent of variation was again relatively mild. Among these 18 sites 28S-Gm789, 28S-Cm3422 and 18S-Cm1303 were differentially methylated by at least 0.05. Interestingly flies grown at 29°C had systematically higher Nm levels while those grown at 18°C tended to have lower Nm levels.

In conclusion, these experiments indicate that Nm on rRNA is quite stable and relatively insensitive to environmental changes. In addition, it appears that the little variations are distinct depending of the stress that was applied.

## Discussion

Along with pseudouridine, Nm is the most abundant chemical modification found on rRNAs. Together with post-translational modifications and paralogs of ribosomal proteins, rRNA modifications contribute to the emerging field of ribosome heterogeneity (45, 46). Several studies show the importance of individual Nm modifications on translation in mammalian cell models as well as on fitness in yeast and archaea (12, 47–49). Nevertheless, much less is known on how differences in Nm between tissues or environmental conditions shape the proteome and therefore the identity of a cell. *Drosophila melanogaster* provides many benefits for studying the role of snoRNAs and of Nm sites on ribosomes because a) it is a multicellular organism and b) has a relatively short lifespan both of which allow to study snoRNA expression and Nm levels in a spatiotemporal manner; c) it presents limited snoRNA gene duplications in comparison to mammals, which makes snoRNA knockouts functional (50) and finally d) a multitude of genetic tools exist for spatiotemporal re-activation of silenced genes.

To fully profit of these aspects, we started by mapping Nm sites on rRNA using two orthogonal techniques: RiboMethSeq and Nanopore DRS. This yielded a list of 61 Nm sites, 50 of which are also conserved in human. In parallel, we sequenced snoRNAs using TGIRT-seq, a customized NGS-based library particularly fit for small and complex small RNAs (29). This data allowed us to faithfully validate expressed snoRNAs, to curate their coordinates, to uniformly predict rRNA targets and to flag snoRNAs that come from likely wrong annotations. The set of confident C/D box snoRNAs were completely renamed to resemble the human nomenclature. Finally, we saw that Nm levels are overall very stable in response to environmental stresses with a few exceptions.

First, we applied a series of filters to RMS data to identify 61 Nm sites in *D. melanogaster*. These briefly consisted into filtering MethScores that are higher than a non-methylated control, up-ranking conserved sites, matching snoRNAs and being responsive to Fib. This conservative approach was completed by the validation of a large majority of sites by Nanopore. Although we found that the potency of Nanopore DRS for *de novo* discovery of Nm sites was limited, we showed nevertheless that Nanopore was an excellent tool for validating RMS hits. Independently of the method, we observed a highly variable range of response to Fib KD. Interestingly, this did not correlate with snoRNA expression nor number of snoRNA isoforms matching each site (**Supplementary Figure 7**). We tried to address more parameters that could explain this heterogeneous response to Fib KD without any success, such as the conservation of Nm sites, their secondary structure context or even the order in which they are methylated during ribogenesis. Although the order of Nm deposition has been divided into early (co-transcriptional) and later times (nucleolar) in yeast, the actual sequence of deposition remains approximate (51). In addition to this existing complexity, Nm deposition on specific sites depends on nucleolar factors beyond the core snoRNP (52). In fact, the snoRNA:rRNA association does not serve only the purpose of depositing Nm. It is essential for bringing together distant rRNA sequences and give to snoRNP complexes the time to fold rRNA in the right conformation (2). Therefore, as the knowledge of function of individual Nm sites *per se* is very limited (12, 47), our map of Nm sites in *D. melanogaster* along with that of other model organisms is setting the base for future functional studies (**Supplementary Table 4**).

The only way currently to study the function of individual Nm sites is through the depletion of their corresponding snoRNAs. Since the annotation of C/D box snoRNAs and their rRNA targets in *D. melanogaster* were outdated we curated the coordinates of expressed snoRNAs and improved considerably rRNA target predictions in a traceable manner. We used only snoScan to predict canonical rRNA targets, yet we find it to be the most complete tool as it covers the limitations of other tools: snoReport looks for C/D box motifs in stringent positions in the transcript, PLEXY requires knowing aforehand the location and sequence of D/D’ boxes and snoGlobe seeks only for anti-sense element complementarity, but not for C/D boxes (39, 53, 54). Here, we propose *bona fide* annotations for 106 C/D box snoRNAs, a quarter of which were predicted to be an exclusive match for 27 individual Nm sites. This feature is particularly interesting for generating single-snoRNA mutants and studying the demethylation of single Nm sites. The 37 previously annotated, but not expressed snoRNAs are perhaps not expressed in the tissues and developmental stages that we examined or not covered enough by our sequencing. Nevertheless, we favour the hypothesis that 25 out of them are falsely annotated as genes since there is no evidence of expression in literature. Instead they could be classified as *pseudogene* (**Figure 3A**). This is supported by the fact that host gene expression is a positive indicator of intronic snoRNA expression, which is the opposite in the case of 34 out of the 37 non expressed snoRNAs whose respective host gene is expressed (50). Finally, our *de novo* search for non-annotated snoRNAs yielded only one hit, snoCD45, but we cannot exclude that future tools or different parameters could reveal new C/D box snoRNAs in *D. melanogaster*.

The canonical role of snoRNAs is guiding chemical modifications on rRNAs. Based on this, they were named after the main rRNA target-nucleotide in *D. melanogaster*. Because there is growing evidence that snoRNAs are involved in the modification of other RNAs and mechanisms such as splicing (39, 55, 56), we saw the need to propose a new nomenclature. The latter is free of any association to specific targets and only reveals the class it belongs to (in this case C/D box). As our nomenclature is numeric-based, it can easily be extended in the future by any putative snoRNA discoveries.

As part of other modulable features that contribute to ribosome heterogeneity, rRNA modifications are also thought to shape the translatome (57). This specialisation has been observed during development and to a lesser extent in response to environmental cues (58, 59). rRNA modifications are perhaps not surprisingly resistant to the latter (24, 52). Given they are mostly deposited during ribosome biogenesis and that their half-life is in the order of several hours in bacteria and 3-5 days in mammalian cells, a certain degree of ribosomal turnover must occur to observe changes (BioNumbers 108025, 108023, 110053) (44). In addition, there are no enzyme known to erase rRNA modifications. Based on literature, we selected environmental stresses that affect ribosome composition or abundance. So far, heat is the only stress known to affect rRNA modifications, particularly in *T. kodakarensis* where the number of acetylated cytidines increases dramatically when these archaea are grown in 85°C instead of 55°C (49). Other stresses such as acute nutrient deprivation or oxidative stress can affect RNA modifications indirectly by reducing the number of ribosomes or damaging their integrity, respectively (60, 61). With these examples in mind, we tested oxidative, nutrient and heat stresses for 8 to 10 days, a duration that is sufficient for significant renewal of the ribosome pool. The mild difference observed in 6 Nm sites across all stresses could not be linked neither to a change of the corresponding snoRNA expression, nor to the expression of the stand-alone CG8939 (data not shown). This very limited variation of Nm levels indicate that Nm modification machinery is very resistant to ambient stresses in *D. melanogaster*. Yet the significant variation of certain Nm sites we observed might serve as a functional response to adapt to the environment.

In conclusion, our work opens the door to studying the role of rRNA Nm sites at the whole organism level. With a confident map of Nm sites and a better characterised set of matching snoRNAs, it will become feasible to individually manipulate Nm sites and study their function in specific tissues and developmental stages.

## Data availability

Basecalled FAST5 of nanopore direct RNA sequencing runs have been deposited to ENA, under accession code PRJEB45722. Tracks for IGV visualization of the mapped reads (BAM files), predicted rRNA modifications in *D. melanogaster* (based on homology to known human rRNA modifications), *EpiNano* RNA modification scores, as well as the code to perform *EpiNano* score analysis per k-mer and *de novo* discovery of Nm sites, can be found in GitHub (https://github.com/novoalab/Nm_Nanopore_Drosophila).

## Acknowledgements

We thank members of the Roignant, Motorin and Novoa labs for helpful discussion. We acknowledge the support of the MEIC to the EMBL partnership, Centro de Excelencia Severo Ochoa and CERCA Programme / Generalitat de Catalunya.

