## Supplementary Methods and Figures for "Comprehensive map of ribosomal 2′-O-methylation and C/D box snoRNAs in *Drosophila melanogaster*"

#### Pipeline of snoRNA:Nm sites predictions

The 142 corrected fasta files were screened for C/D snoRNA features and for putative Nm targets on rRNA sequences using SnoScan (overall score ≥ 6). Because the flexible cut-off returned between 1 and 52 predictions per snoRNA, we decided to consider the three best scoring ones. Then we introduced a filter based on the candidates that matched the previously identified 62 Nm sites. From the 146 predictions, 131 were further kept based on whether they were the $1^{st}$ best hit or if the $2^{nd}$ best hit was consecutive to another confident Nm site. Indeed, in other organisms, C/D snoRNAs can target two – usually consecutive – Nm targets. Secondary targets were also kept if at least 10 bp of the ASE aligned to rRNA sequences without a single mismatch but including Hoogsteen base pairing. In total, 95 expressed transcripts were confidently predicted to target 119 Nm sites while 12 transcripts were not covered and were therefore ranked as “Low confidence”.

For the remaining predictions of unknown targets, only the first best hit was kept for the predictions targeting unknown Nm sites, unless the second best prediction targeted a confident Nm site, in which case the candidate was already included in the confident Nm site group. After this filter, we checked whether the 27 Nm candidates predicted by us, but also those predicted by their original publications, were conserved or had a Methscore ≥ 0.82. None of them met these criteria, therefore the 7 transcripts that were expressed were labelled as orphan while the remaining 20 were classified with the low confident predictions. Finally, 8 annotated transcripts yielded no prediction at all, 3 of which were expressed and were labelled as “orphan” and 5 which were not expressed and were and classified with the remaining “Low confidence” snoRNAs.

### Supplementary Figure 1

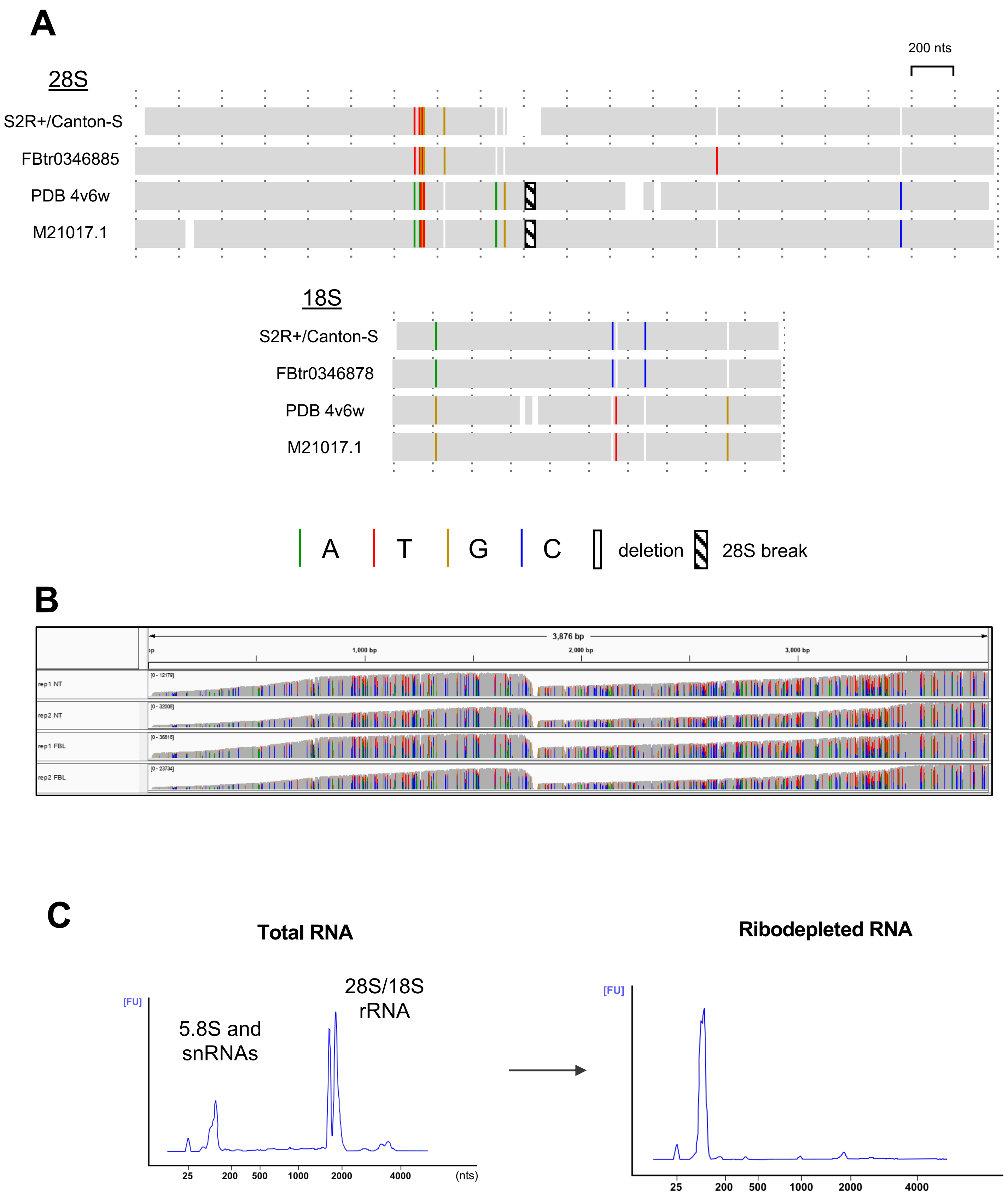


**Supplementary Figure 1**
(A) Summary of sequence differences between main rRNA references and Sanger sequencing of 28S and 18S in both S2R+ and Canton-S flies. (B) Read coverage of native 28S rRNA obtained with Nanopore. (C) RNA profiles before and after ribodepletion (Bioanalyzer and siTools)

### Supplementary Figure 2

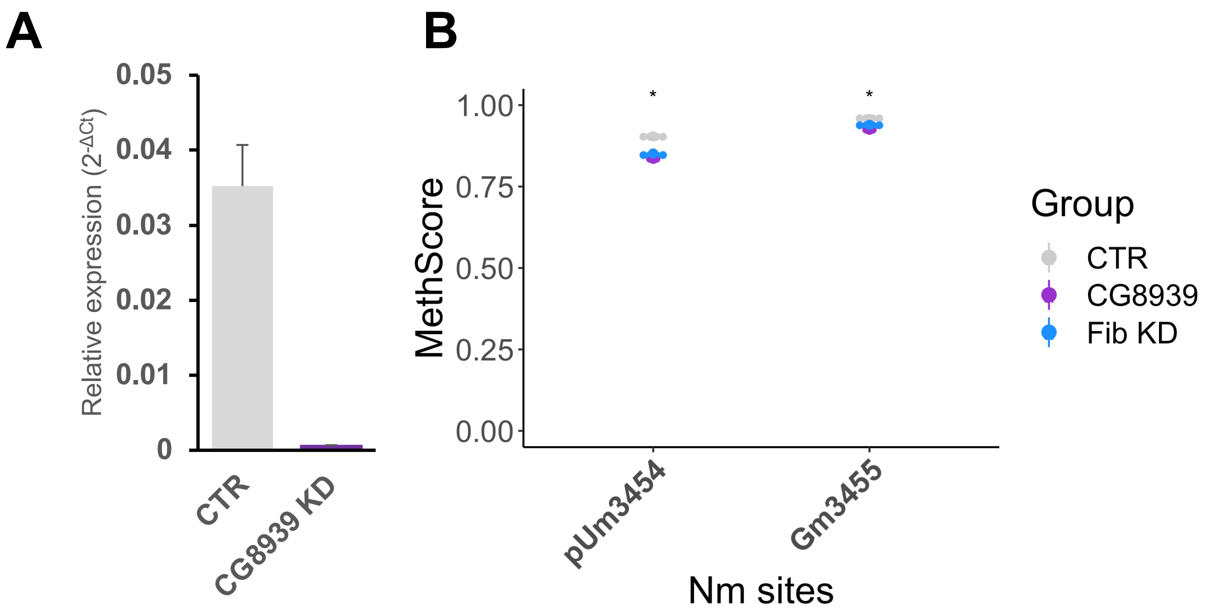


**Supplementary Figure 2**
(A) Efficiency of CG8939 (FTSJ3) knockdown in S2R+ cells, quantified by qPCR (n=3)
(B) Nm levels of Nm sites that vary with CG8939 KD (Kruskal-Wallis test, * p<0.05, n=3)

### Supplementary Figure 3A

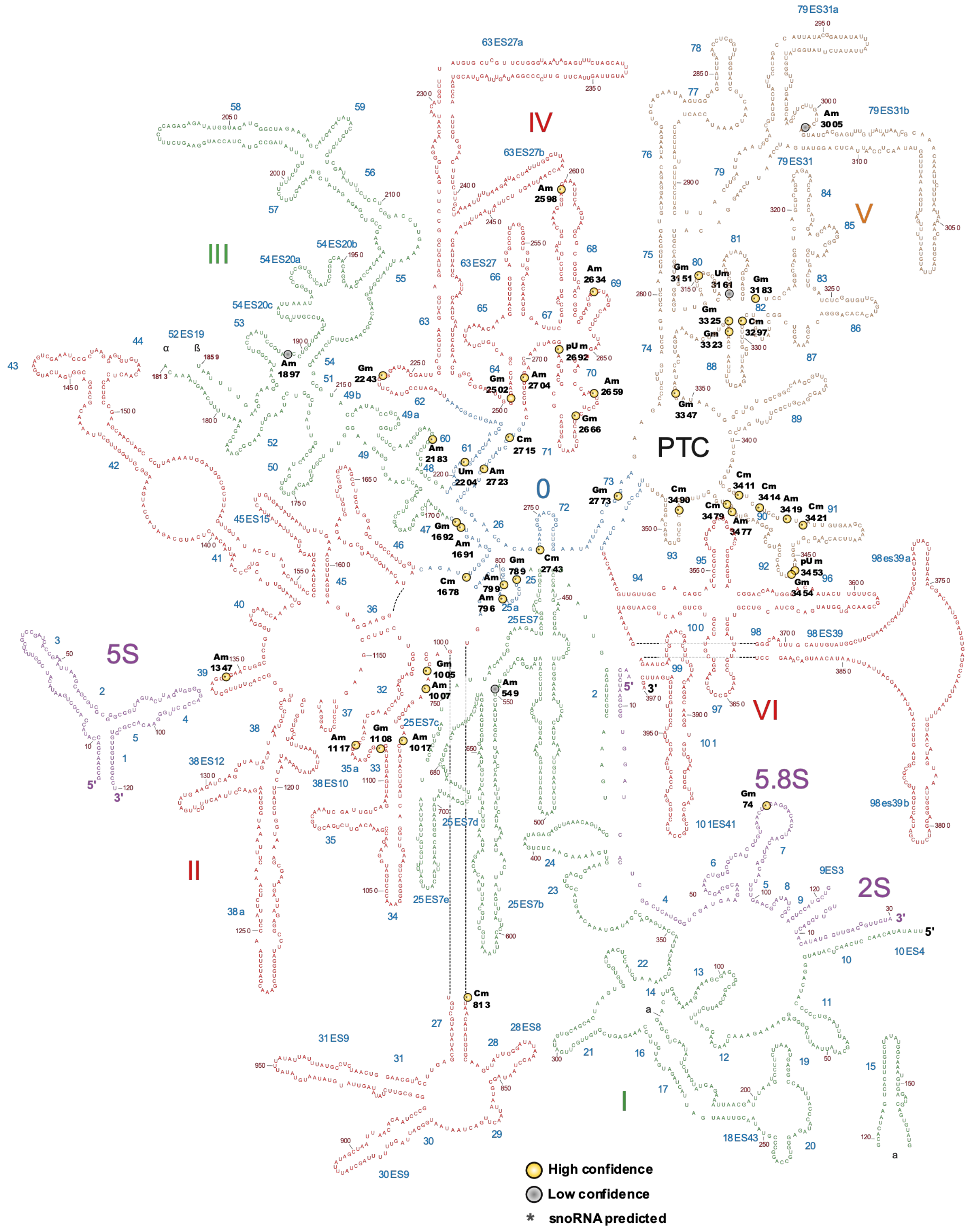


**Supplementary Figure 3**
(A) 2D projection of Large Subunit rRNAs (28S, 5.8S, 2S and 5S) annnotated with high and low confidence Nm sites

### Supplementary Figure 3B

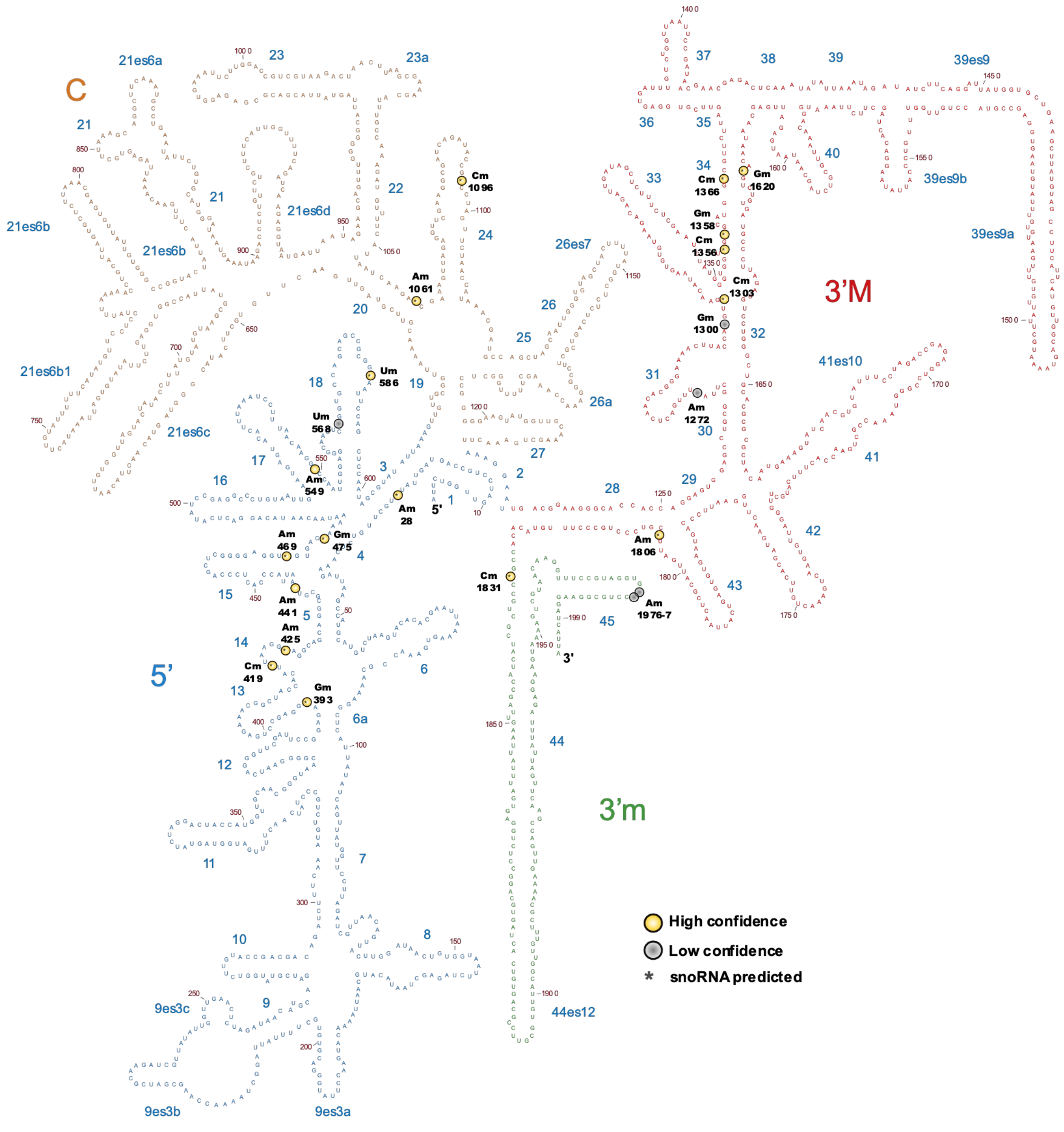


**Supplementary Figure 3 (continuation**

(B) 2D projection of Small Subunit rRNA (18S) annnotated with high and low confidence Nm sites

### Supplementary Figure 4

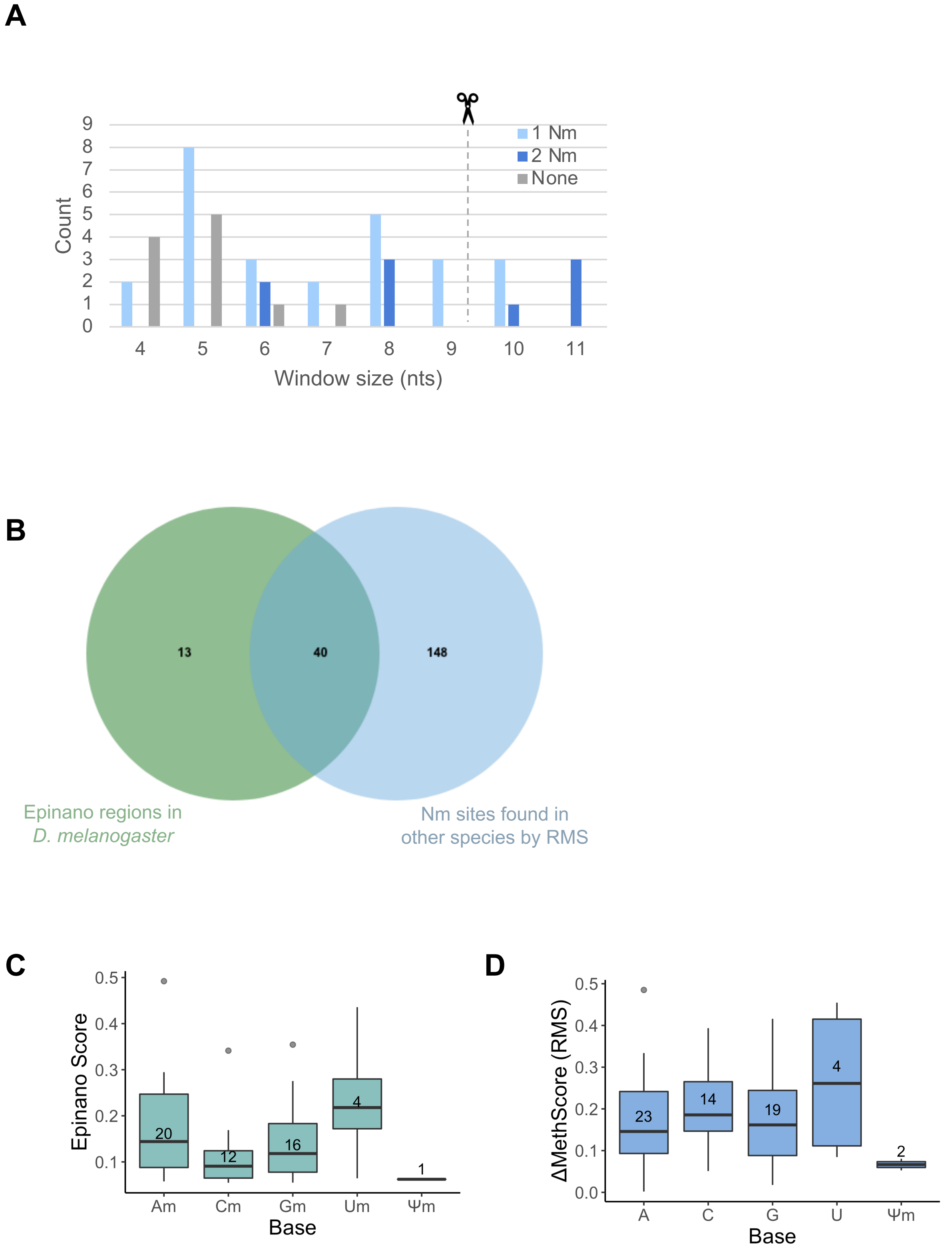


**Supplementary Figure 4**

(A) Size distribution of regions (nts) with an EpiNano score of ≥3x median in both replicates or ≥5x median in one replicate, before splitting large regions into 7mers. Windows are separated in regions that overlap 1Nm, 2 Nm or no Nm sites from the RMS dataset. (B) Overlap of EpiNano 7 mer regions with Nm sites that are found in RMS data from *D. melanogaster*, *H. sapiens*, *S. cerevisiae*, *X. laevis*, *A. thaliana* and *D. discoideum*. (C) Distribution of Epinano Score and of (D) RMS ΔMethScore for each base.

### Supplementary Figure 5

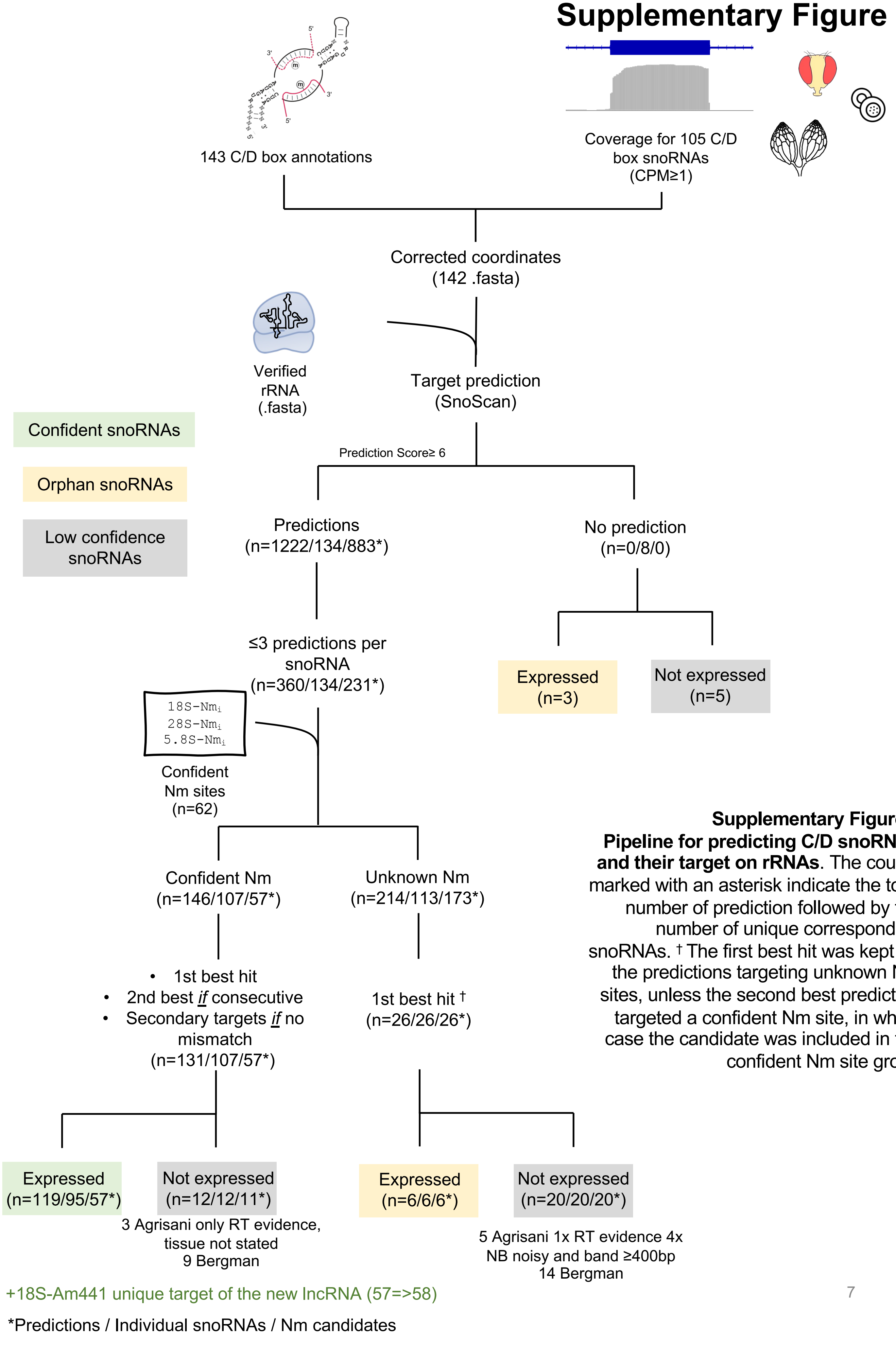


**Supplementary Figure 5**
**Pipeline for predicting C/D snoRNAs and their target on rRNAs**. The counts marked with an asterisk indicate the total number of prediction followed by the number of unique corresponding snoRNAs. † The first best hit was kept for the predictions targeting unknown Nm sites, unless the second best prediction targeted a confident Nm site, in which case the candidate was included in the confident Nm site group

### Supplementary Figure 6

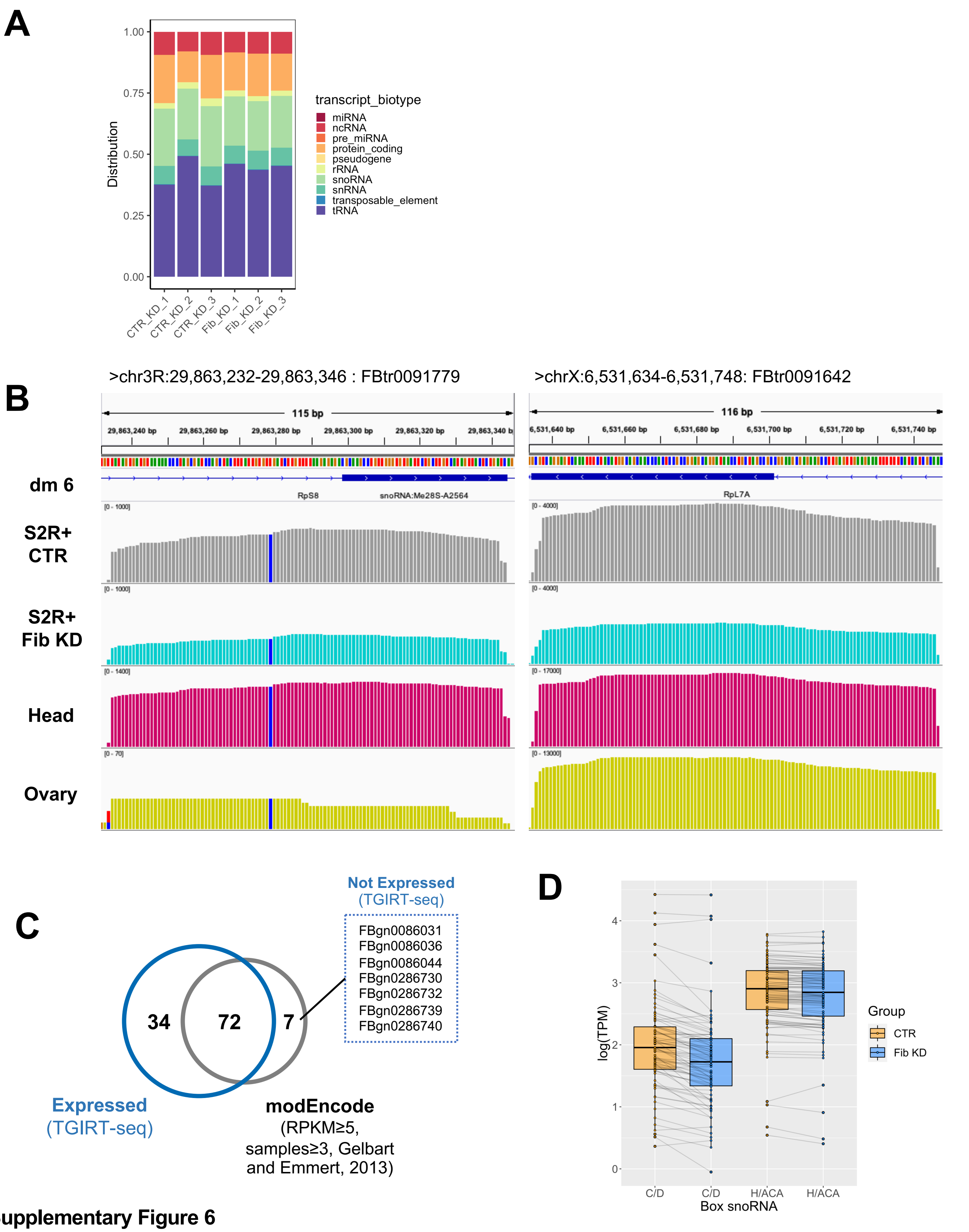


**Supplementary Figure 6**

(A) TPM distribution across different RNA classes in TGIRT-seq of S2R+ cells. (B) Examples of discrepant genomic coordinates between dm6 (BDGP 6.28) annotations and read coverage across S2R+ CTR, S2R+ Fib KD, head and ovary samples. (C) Overlap of expressed genes detected by TGIRT-seq in this study and by classic polyA sequencing across different tissues and developmental stages (modENCODE). (D) Average expression levels of C/D and H/ACA box snoRNAs in CTR and Fib KD S2R+ cells (n=3)

### Supplementary Figure 7

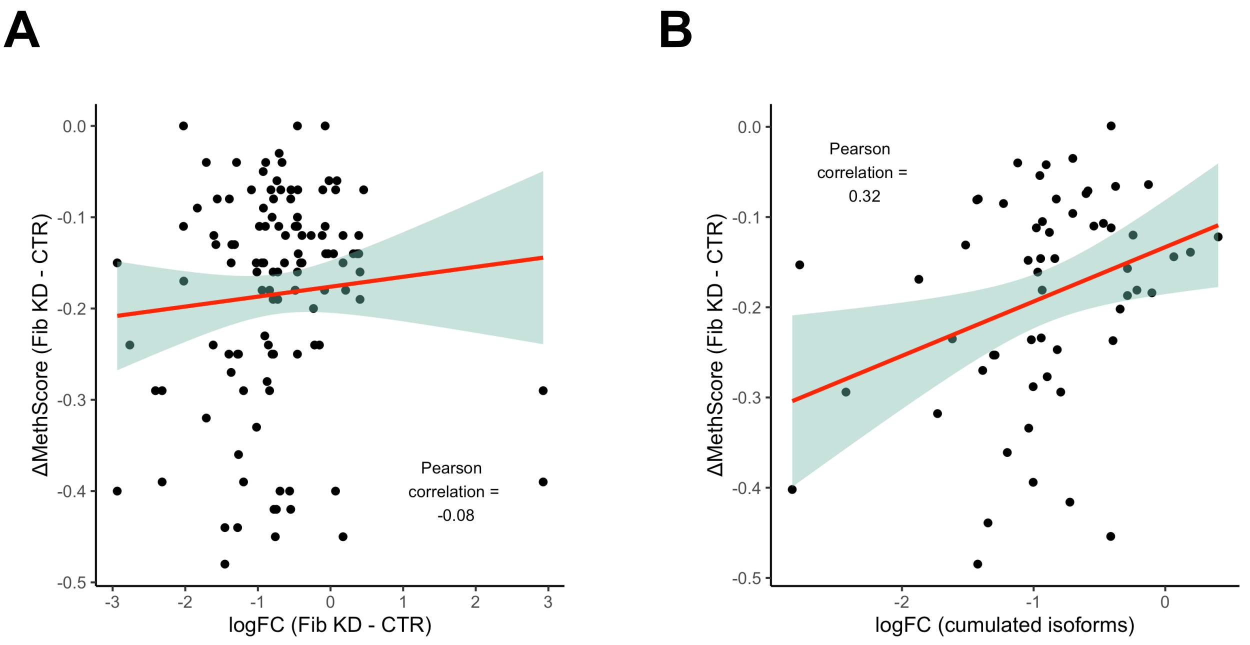


**Supplementary Figure 7**
(A) Correlation of differential methylation (ΔMethscores) and differential expression of matching C/D box snoRNAs (logFC of normalised reads) after comparing Fibrillarin KD to control conditions in S2R+ cells.
(B) Same plot after cumulating normalised reads of snoRNA isoforms that target the same Nm site.
Points were fitted with a linear regression (red line).
